# Allosteric regulation of Glutamate dehydrogenase deamination activity

**DOI:** 10.1101/786954

**Authors:** Mubasher Rashid, Soumen Bera, Alexander B. Medvinsky, Gui-Quan Sun, Bai-Lian Li, Claudia Acquisti, Adnan Sljoka, Amit Chakraborty

## Abstract

Glutamate dehydrogenase (GDH) is a key enzyme interlinking carbon and nitrogen metabolism. Recent discoveries of the GDH specific role in breast cancer, hyperinsulinism/hyperammonemia syndrome, and neurodegenerative diseases have reinvigorated interest on GDH regulation, which remains poorly understood despite extensive and long standing studies. Notwithstanding the growing evidence of the complexity of the allosteric network behind GDH regulation, a clear understanding of the enzyme dynamics involved in the process is limited to the activator Adenosine diphosphate (ADP) and the inhibitor Guanosine triphosphate (GTP). Presently, the identification of further factors involved in the allosteric network is paramount to deepen our understanding of the complex dynamics that regulate GDH enzymatic activity. Combining structural analyses of cryo-electron microscopy data with molecular dynamic simulations, here we show that the cofactor NADH is a key player in the GDH regulation process. Our structural analysis indicates that, binding to the regulatory sites in proximity of the antenna region, NADH acts as a positive allosteric modulator by enhancing both the affinity of GTP binding and inhibition of GDH catalytic activity. We further show that the binding of GTP to the NADH bound GDH produces a local conformational rearrangement that triggers an anticlockwise rotational motion of interlinked alpha-helices with specific tilted helical extension. This structural transition is a fundamental switch in the GDH enzymatic activity. It introduces a torsional stress, and the associated rotational shift in the Rossmann fold closes the catalytic cleft with consequent inhibition of the deamination process. These results shed new light on GDH regulation and may lay new foundation in the design of allosteric agents.

## Introduction

Protein conformational dynamics are mainly regulated by changes in structural flexibilities. However the impact of the associated conformational rearrangements of amino acid residues on protein functions, ligand-bindings, and allosteric communications still remains poorly understood (Beckett, 2009; Nussinov et al., 2013; Wodak et al., 2019). Comparative structural studies have shown the critical role of conformational rearrangements upon ligand binding to allosteric proteins (Teague, 2003; Tukuriki and Tawfik, 2009; Beckett, 2009; Ben-David et al., 2019). These ongoing protein dynamics are highly complex, with several allosteric inhibitors potentially able to act at a large distance from the core catalytic domains and to inhibit the enzymatic activity in cooperation with a variety of cofactors (Kim et al., 2017; Nussinov et al., 2013). Here we investigate the allosteric regulation of the mammalian glutamate dehydrogenase GDH, and show how the Guanosine triphosphate (GTP) binding establishes rigidity-based allosteric communication with the regulatory coenzyme NADH which then synergistically inhibits the oxidative deamination activity of GDH by stabilizing its closed conformation.

GDH provides an essential link between carbon and nitrogen metabolisms (Rife and Cleland, 1980; Bera et al., 2016). Despite extensive studies for the last 40 years, its regulation remains elusive (Stanley et al., 1998; Li et al., 2011). Recent discoveries of GDH specific role in breast cancer (Spinelli et al., 2017), hyperinsulinism/hyperammonemia (HI/HA) syndrome (Grimaldi et al., 2017), and neurodegenerative diseases (Kim et al., 2016) has reinvigorated interest on GDH regulation and given new momentum to the field. While the molecular structure of GDH is well resolved, the underlying dynamics of GDH regulatory system which involves several allosteric modulators remains poorly understood. Important input has come from the genetic variation associated with the hyperinsulinism/hyperammonemia (HI/HA) syndrome in the human population. It was shown that most mutations are located near the GTP binding sites or in the adjacent antenna region (Stanley et al., 1998; Stanley et al., 2011). Mutant GDH from patient lymphoblasts and in the mutant GDH expressed in *E.coli* showed almost complete unresponsiveness of GDH to GTP inhibition, normally reduced from 1.3 to 10-fold compared to the wild type (Allen et al., 2004).The molecular basis of the GTP inhibition and the associated GTP insensitivity are one of the key remaining open questions of the GDH regulatory system.

The role of the cofactor NADH in the regulation of GDH activity has emerged only very recently. It was previously observed that in absence of NADH, GTP binds only weakly to GDH. Single-particle cyro-electron microscopy (cyro-EM) studies have further revealed that the GTP binding affinity is increased in presence of NADH at the regulatory site, and GTP synergistically displaces the complex towards the closed conformational state (Borgnia et al., 2016). However, the underlying mechanism explaining this effect has not been previously elucidated. A key problem is that these combinations are not amenable to crystallization due to the fleeting nature of the structures involved. Although there are several solved crystallographic structures of GDH complexes, these are either crystalized with cofactors and nucleotides or predominantly in closed conformation only. Using the available cyro-EM data of (i) the apo enzyme GDH without bound substrates (PDB id: 3jcz), (ii) the open form of NADH- and GTP bound GDH system (PDB id: 3jd3), and (iii) the closed form of NADH- and GTP boundbound GDH system (PDB id: 3jd4), here we have focused on describing and mapping out whole allosteric network behind GDH regulation. Our computational analysis and simulations have shown that the cofactor NADH is indeed a key player in the allosteric modulation, where it acts as positive allosteric modulator. The results presented in this study indicate that GTP activates a triangular allosteric network interlinking distant GTP binding sites, regulatory NADH, and catalytic sites. This network controls the Nucleotide-binding domain motion that eventually blocks the catalytic cleft in the GTP-induced inhibition dynamics. Moreover, using *in silico* dominant mutational analysis, we show that impeding the allostery network leads to GTP-insensitivity, resulting in GDH over-activity.

## Results and Discussion

### A large conformational difference between open and closed GDH system

Cyro-EM GDH complex structure exists in open and closed conformations. Both forms have bound cofactor NADH and the inhibitor GTP having substantially different conformations. While the closed form is relatively well resolved and detailed structural knowledge has been accumulated during last two decades, the structure of the open form is not well understood. This is problematic because allosteric regulation is primarily driven by the inherent differences between the open and closed forms. Using a structural superimposition with RMSD fingerprint of individual residues, we have focused on the detail characterization of the differences between the open and closed structures. GDH has a homohexameric structure composed of a trimer of dimers, where each monomer consists of three identified domains (Banerjee et al., 2003): (i) Catalytic domain, (ii) Nucleotide-binding domain (NBD), and the (iii) Antennae. Borgnia et al. (2016) solved the 3D cyro-EM structure of open and closed forms of GDH bound with both NADH and GTP. Within these structures, the catalytic domain is located near the dimer interface. Moreover, in both the open and closed states, adenosine and nicotinamide moieties of the cofactor NADH are present in nearly the same orientation. The Nucleotide-binding domain (NBD) contains the *βαβ* Rossmann fold that undergoes a large conformational change during the transition. We have focused on these particular domains and investigated in detail the local architecture of the catalytic cleft by measuring the distances of all the AA residues. We can clearly associate the closure of the catalytic cleft with changes in the distances between the following pairs of amino acid residues: Asn254-Thr171, Asn349-Asp168, and Ser276-Lys134. These pairwise distances are reduced at least 2-fold in the transition from the open to the closed form (Fig.1B).

**Fig. 1.**
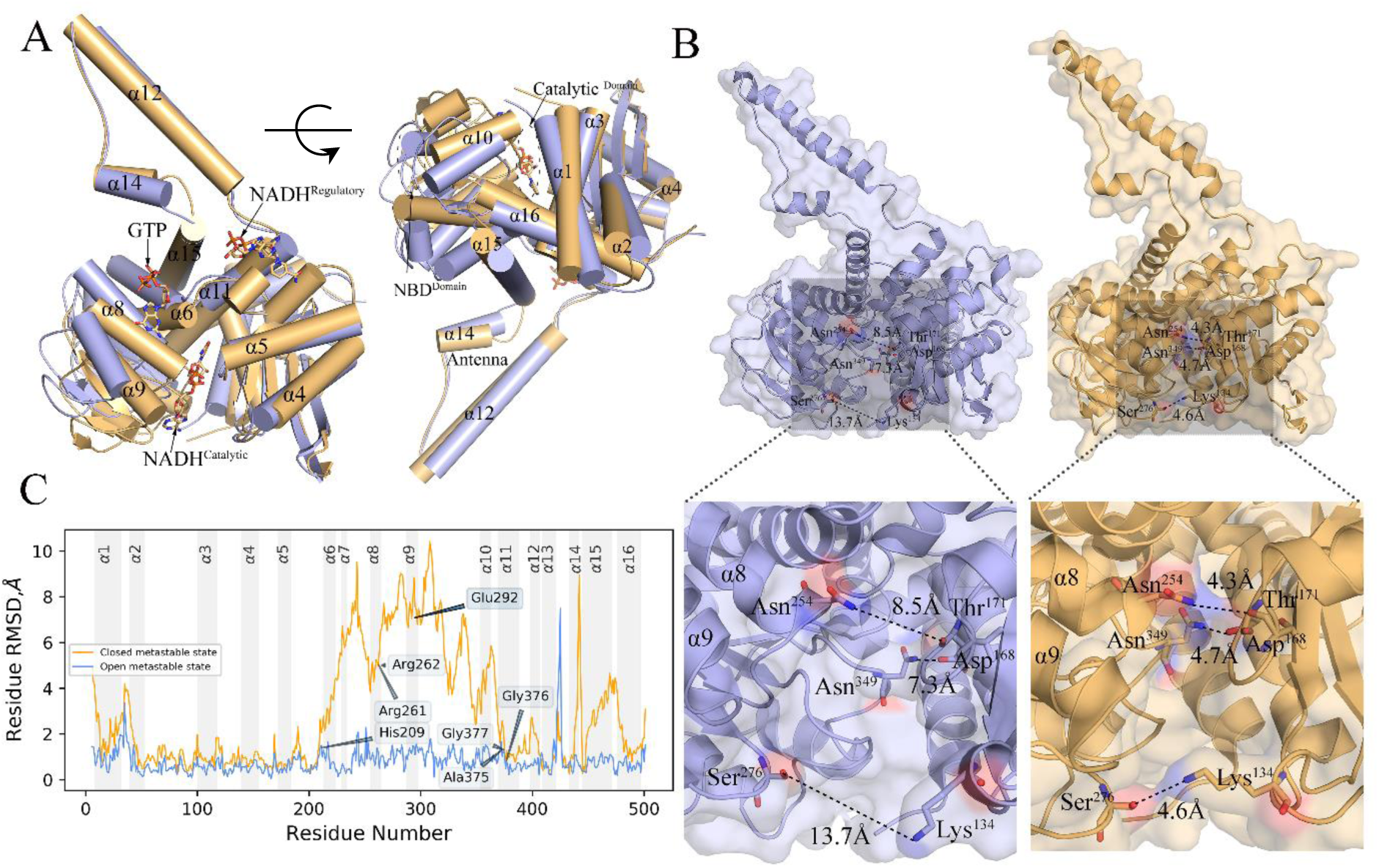
Conformational differences between the open-and closed state of the GDH regulatory system, indicated by light blue and light orange color respectively. Two structures are superimposed using atoms and α-helices. (A) 3D side view of opposite sides of α-helices and their transitional and rotational shifts. Most α-helical shifts are local at NDB domain in-between the GTP-binding sites and NADH-boundbound catalytic sites. (B) Local distance between surface residues within the GDH-catalytic cavity significantly reduces at the closed state relative to its open conformer. (C) The root means square deviations (RMSDs) of C*α* atoms for each residue conveniently differentiate the two conformational states, with substantial conformational changes in the helices VI to XII and the antennae regions.

It was previously noted that within the catalytic cavity, both the adenosine and nicotinamide moieties of bound-NADH keep the same orientation irrespective of conformational forms (Borgnia et al., 2016). Our detailed examination of this domain shows that eight hydrogen bonds keep the NADH bound in the catalytic pocket in both forms (**Supplementary table 2**). Comparison of the closed and open structures points to a critical shift in the positions of Ans349 and Ser276 residues involved in the formation of H-bonds. In the closed form Ans349 is the closest residue to NADH (2.45), while in the open form Ser276 is the closest residue (2.47) forming H-bond to the catalytic NADH. Furthermore, we find that the composition of AA residues that built-up the catalytic cleft differs significantly between the two forms. Asn374, Ala348, Ala326, Val255, Gly253, Gly251, and Thr215 contribute the most to make-up the catalytic cleft in the open form, while Asn374, Ala348, Ala326, Ala325, Gly256, Val255, Gly251, Thr215, Asp168, Pro167 are the primary adjacent residues in the closed form (**Supplementary figure 3**). To determine whether the GDH-NADH interactions in the catalytic cleft change during the transition, we measure local interface area between NADH and GDH. We find that the interface area is reduced during transition from open to closed form, indicating a reduction of intrinsic disorder with decreasing interactions (**Supplementary figure 4**).

**Fig. 2.**
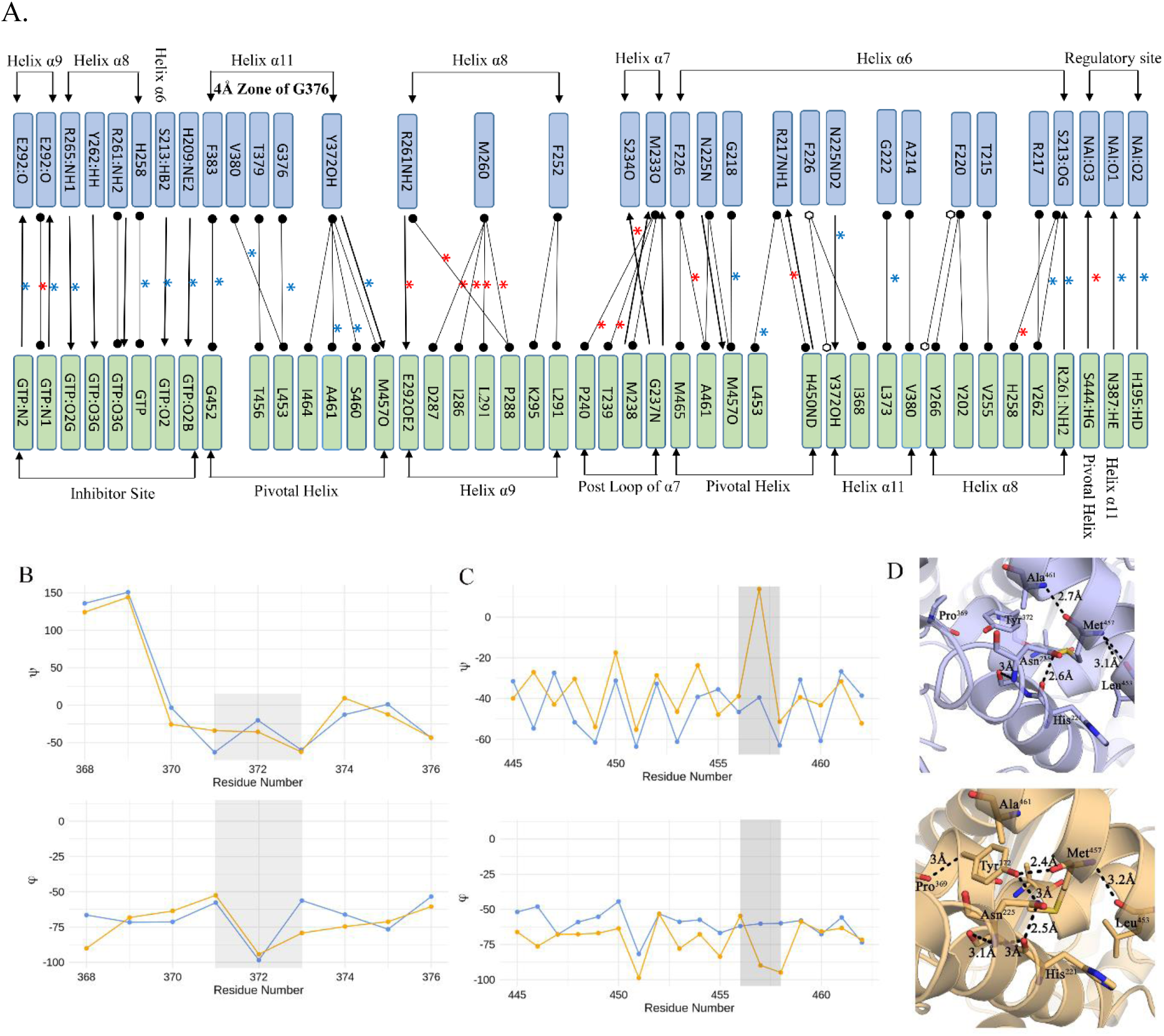
Specific interhelical interactions that are changed due to conformational rearrangement of helices during GTP-triggered transition from open (blue) to closed (yellow) GDH state. (A) (*) represents absent interaction in the open form and (*) indicates absent in the closed form, while hydrogen-bond, van der Waals, and π-π interactions are notated as →, 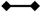 and 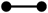. (B) and (C) show the transitional changes in dihedral angles of residue Y372 of *α*11 and M457 of the pivotal helix. (D) Structural snapshots that represent the interhelical interactions between pivotal helix, *α*11 and *α*6 establish in the closed form through H-bond interactions M457 (pivotal helix), Y372 (*α*11) and H221 (*α*6)

**Fig. 3.**
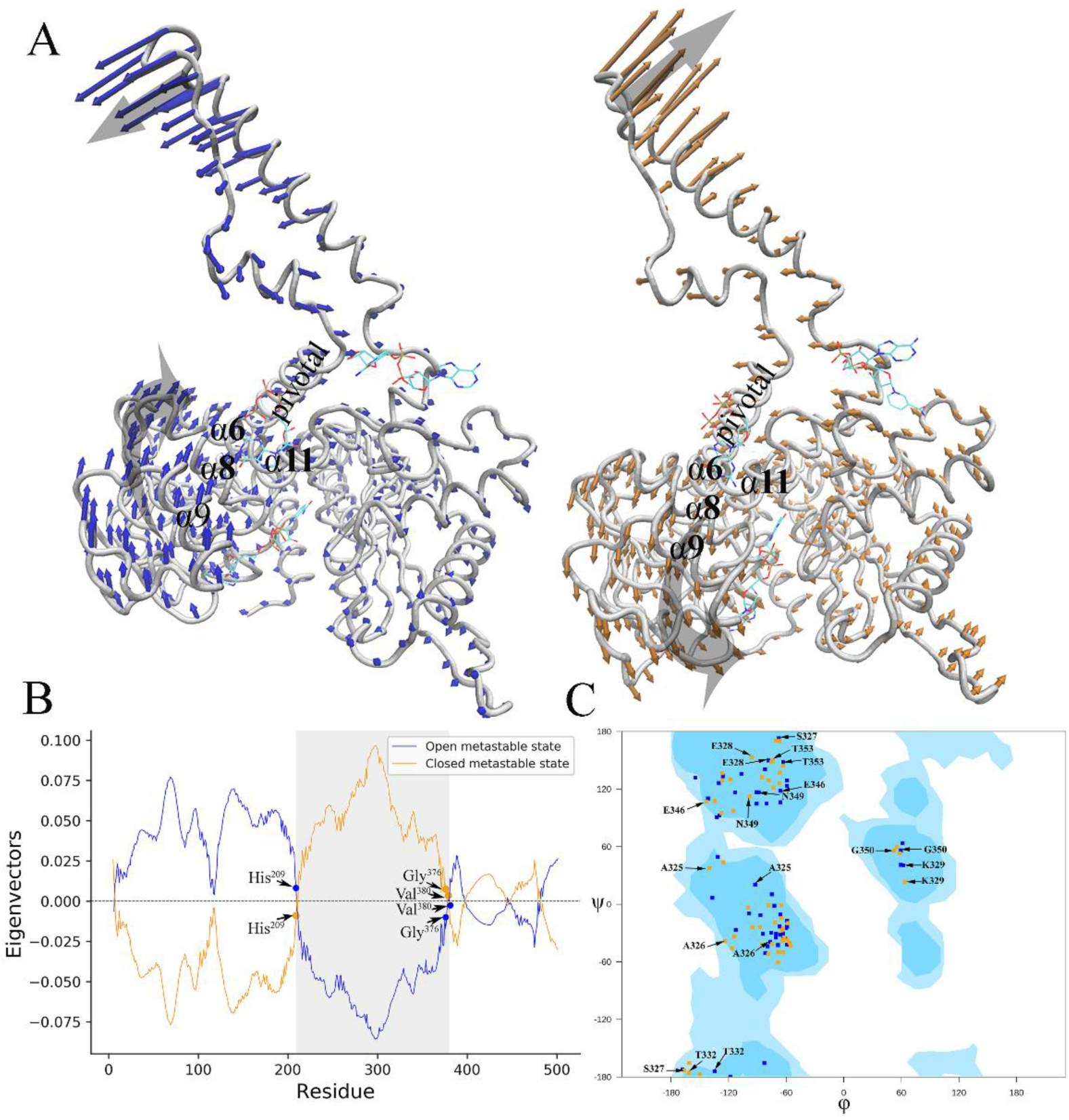
Relative directions and magnitudes of flexibility of open -and closed state of GDH system from low-frequency mode of a three-dimensional Elastic Network Model. (A) Opposite domain motions can be seen near to the equilibrium for the open (light blue) and closed (light orange) conformers. (B) The region around the AA residue 200-400 involving the helices VI to XII attained the picks of opposite motions and highest RMSD differences between the two states. (C) Ramachandran plot of the Rossmann fold AA residues that were superimposed on the contour plot generated with all phi-psi AA confirmations of 1000 frames generated by MD simulation. It indicates rotational shift of the residues as the open conformer moves to closed form.

**Fig. 4.**
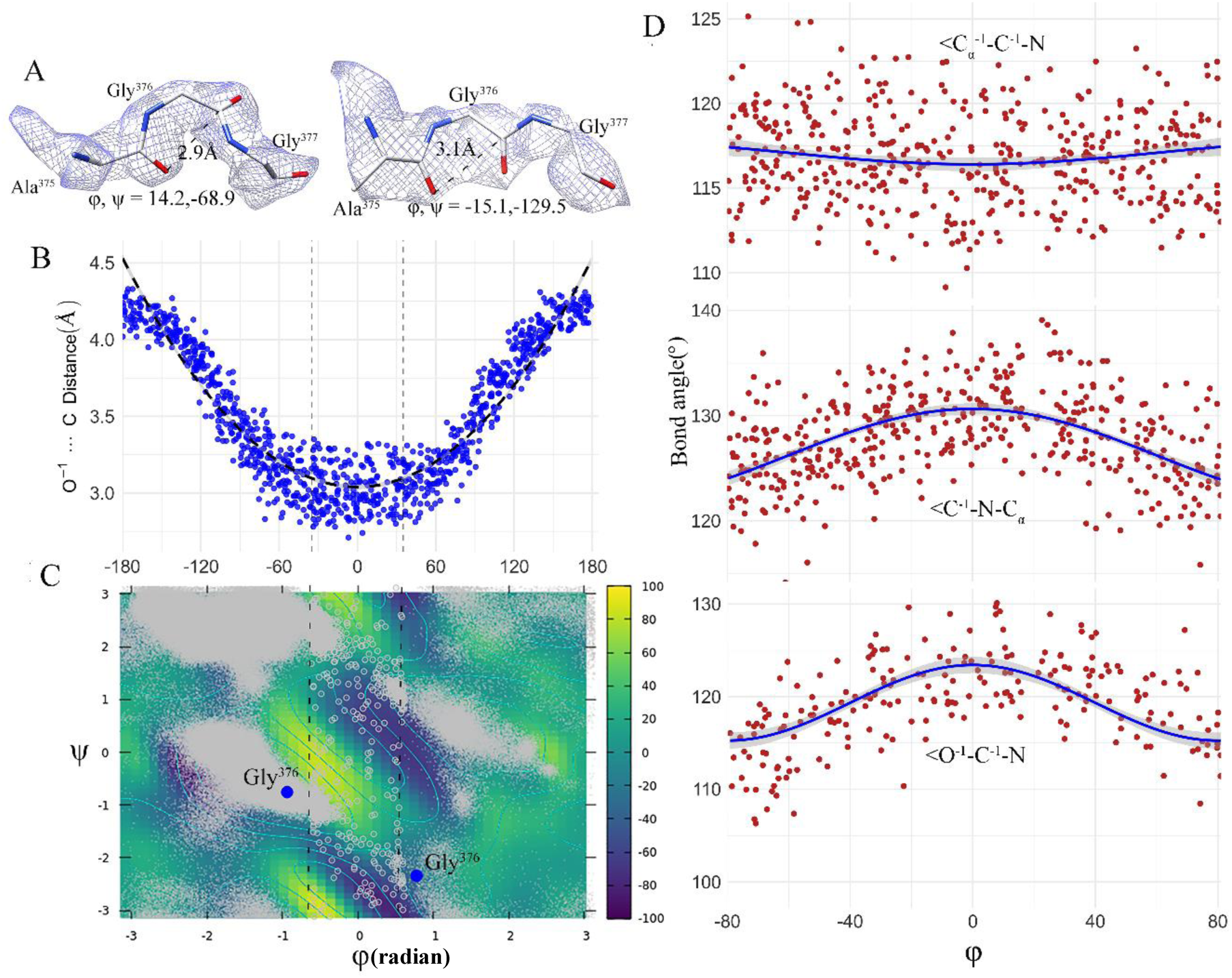
Transition residues for conformational changes from GDH open to closed state. (A) cryo-EM map evidence for transient occurrences of Gly 376 in the transition zone (−35° ≤ *φ* ≤ + 35°) which is a disallowed region in the Ramachandran phi-psi plot of protein backbone conformations. These peptide conformations are the randomly chosen snapshot representations at 2.49ns and 6.5ns from the 1000 frames generated at every 10 picoseconds by MD simulations of open conformer that ran for 10 nanoseconds using the force field CHARMM36. (B) Distortions of the distance O^-1^… C with respect to changes of φ along the MD trajectory. MD predicted data points (blue circle) are well-fitted with a quadratic function of φ: y= (5E − 05) *φ*^2^ + 0.001 *φ* + 3.0397 (*R*^2^ = 0.8813). (C) A Ramachandran plot with energy contours for the tripeptide Ala375-Gly376-Gly377 calculated using adaptive biasing force methods. It used residues from 1000 frames generated by MD simulations. Small grey dots refer to non-transient residues sitting in the allowable regions of Ramachnadran plot and grey circle refers to transient residues located at the transition zone. It further showed that Gly 376 (blue solid circle) crossed the high energy barrier near to *φ* ≈° through the low energy passes at Ψ ≈ −90°. (D) Systematic distortions of the bond-angles with respect to φ. MD predicted data points are smoothen by a cosine function: Y = *I*(cos(*φ* * *pi* / 120)).

**Fig. 5.**
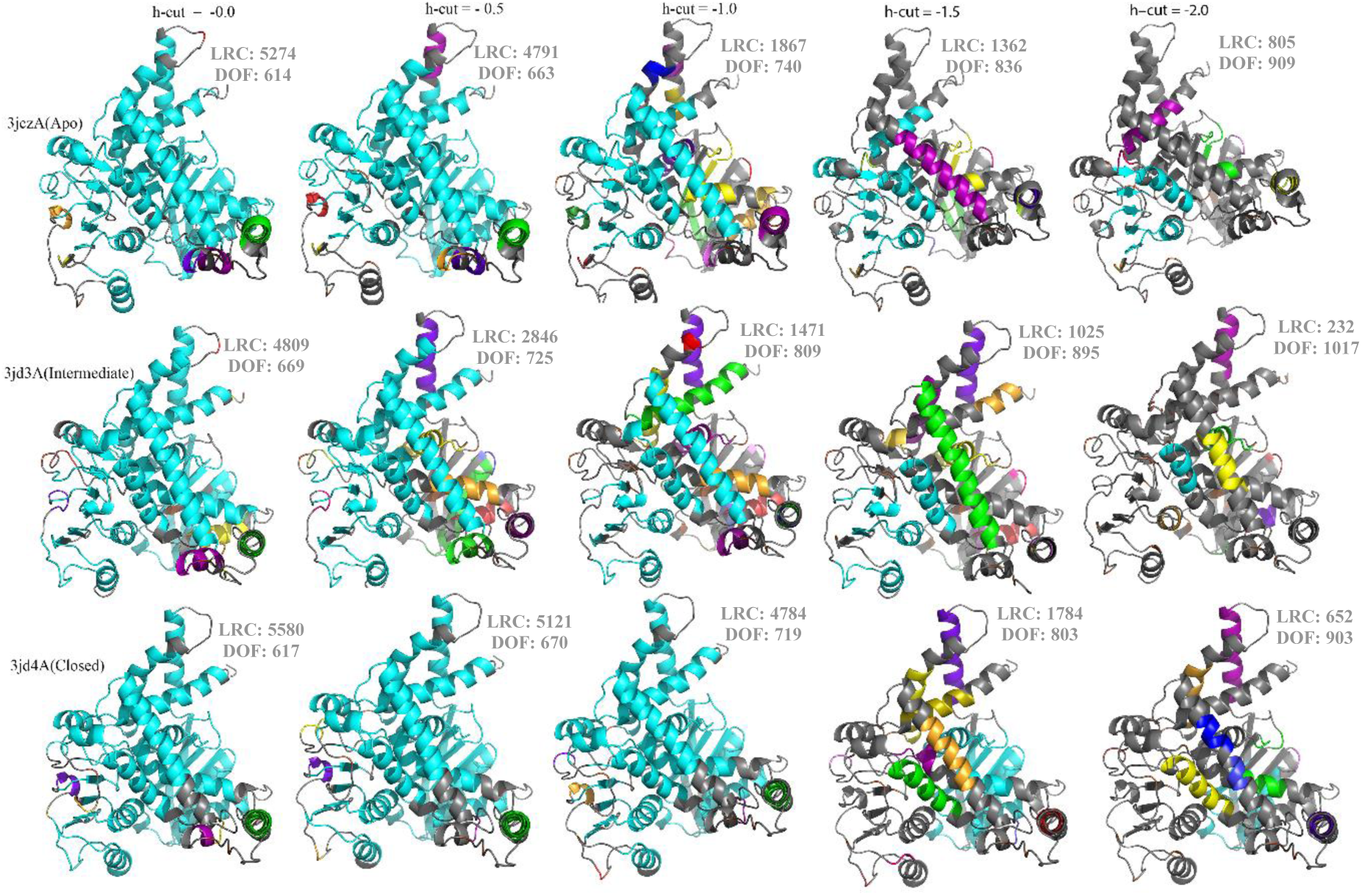
Rigid cluster decomposition of GDH system with various hydrogen-bond energy cut-offs. The largest rigid cluster (LRC) is represented by cyan color and its size is progressively reduced as H-bond energy cut-off decreases from 0 to -2.0 kcal/mol. H-bond cut-off -0.5 kcal/mol is considered as acceptable cut-off that indicated by the H-bond dilution plot.

At the regulatory sites in both structures which are located at the vicinity of “antennae”, the cofactor NADH is known to be bound as an inhibitor and then mutually facilitates inhibition of GDH deamination activity together with the GTP bound at a distant site. Borgnia et al. (2016) observed that Adenine moiety of NADH has identical binding pocket as ADP, a potent activator of GDH and its conformations remain unaltered between the two conformers. However, nicotinamide portion of NADH has different conformations between open and closed state. It was further reported that His209, which undergoes large movement, plays critical role in NADH binding and stabilization when GDH system moves towards its closed form. Our distance calculations perfectly match this observation that the alpha-phosphate of NADH is ∼4.4 Å away from His209 in the open form, and is more than 10.5 Å away in the closed form. Furthermore, NADH binding engages other interactions and has altered adjacent residual compositions when open structure transits to closed form (**Supplementary figure 2 and Supplementary table 3**).

GTP binding site is located next to pivot helix at the base of antenna in the both the conformers. We find that the GTP-GDH local interface surface area increases (∼48 Å^2^) while it moves towards the closed state (**Supplementary figure 3**), indicating much closer interactions. These increase interactions we observe in the formation of three H-bonds between NADH and His209, Ser213, and Glu292 in the closed state (**Supplementary table 4**), which explain the stabilization of the GTP binding. Interestingly, we note that GTP-GDH interactions are strongly modulated by regulatory NADH with effects on the GTP binding affinity. For *in silico* affinity calculation, we use CSM-lig machine-learning method (Pires and Ascher 2016). In presence of NADH at regulatory sites, our calculation shows that GTP binding affinity is much higher (∼2 Kcal/mol) with more close interactions through 2 additional H-bonding. This close interaction with total 7 H-bonding explains the higher affinity of GTP. We further observe NADH-triggered local conformational rearrangement through inclusion and exclusion of H-bonding. Two closest H-bond interactions between the residue Arg261 and GTP (2.71 Å and 2.74 Å) in absence of regulatory NADH have completely been lost when NADH is binded at the regulatory site, while the AA residues Tyr262 (2.35 Å), His209 (2.36 Å), and Glu292 (2.72 Å) forms strong H-bonds to boundbound GTP in the closed conformational state in presence of regulatory NADH.

We calculated the root means square deviations (RMSDs) of C *α* atoms for each residue for determining interhelical differences between the two conformational states. This comparison shows that the helices I to V form a stable helical bundle core, with minimal structural changes. In contrast, the substantial conformational changes are noted in the helices VI to XII and in the antennae regions (Fig.1A,C). Correlation coefficient between the two sets of RMSDs for I to V is 0.796, indicating strong similarities. Substantial differences in helices VI to XII are signified by the correlation coefficient 0.254 (**Details in Supplementary figure 1 and Supplementary table 1**). These differences are resulted in the rotational motions of the helices, which are indicated in our measurements of helical rotational angles with respect to the apo structure. We find that the highest rotational shift of the helix X (∼ 20.1^0^) and XIV (∼17.8^0^) with respect to corresponding apo structure (**Supplementary figure 4**). The associated rotational directions show anti-clockwise movement of most helices VI to XII and the XIV in the closed state, indicating inclusion and exclusion of some specific interhelical interactions during the transition from open to closed state.

### Specific interhelical interactions are triggered by GTP binding

With the residue interaction networks (RINs) approach (Martin et al., 2011), we examined interhelical interactions and identified specific residue-residue interactions between the helices by comparing the open and closed structures. RIN shows significant changes in the interactions among the helices VI-XII when the GDH transitions from open to closed form. To probe this further, we computationally calculated the differences of ^13^C, ^15^N, and ^1^H chemical shifts (Δ*δ*) using a combined approach of machine learning and sequence alignment (Han et al., 2011). Chemical shifts were calculated for all residues laying on the helices with substantial rotations (**Supplementary figure 9**). Although average differences in ^13^C Δ*δ* with respect to the apo structure for open and closed form (i.e., 0.80 and 0.75 p.p.m) are not significant, many residues of pivotal helix, *α*9, *α*10, and *α*11 show a large differences in Δ*δ*, indicating significant conformational rearrangement.

These calculations show that pivotal helix has retained the maximum number of interactions with other helices in both the forms. The most specific molecular interactions that are formed at the closed conformational state but lost in the open state are between the helix and pivotal helix, including five Van Der Waals interactions and one strong H-bond interactions between Y372 and M457 (Fig.2A, B, C). All these interactions occur within the 4.0 Å neighbourhood of GTP binding site. This reinforces that H-bond contact establishes a link between the pivotal helix,, and via the Y372 that has H-bond contact with N225 of (Fig.2D). Loss of these H-bonds in the open conformation is reflected in ^15^N –^1^H plot of the chemical shifts, in which all the three residues show significant shift in the ^15^N-^1^H space (**Supplementary figure 9**). Additionally, residue N387 forms a H-bond with the regulatory NADH at closed state and thereby it provides a potential allosteric link between GTP binding site and NADH at regulatory site. Formation of these interactions underlies the large conformational differences observed between the open and closed form.

To examine the underlying dynamics of the conformational changes, we analyzed low-frequency motions using elastic network models (ENMs) in DynOmics webserver (Li et al., 2017) that integrates widely used Gaussian network model (GNM) and Anisotropic network model (ANM). Among several independent low-frequency normal modes, we find that the mode 2 convincingly differentiates the domain motions between the open and closed forms (Fig.3A). At the onset of transitions, anti-clockwise helical motions are noted with the pivot helix rotated about 5.7^0^ relative to the apo state. The opposite domain motions along the normal mode of the two metastable states indicates some hinge-bending motions that are important for the transitions. It shows the region around the residue 200-400 involving the helices VI to XII, which attained the picks of the opposite motions and highest RMSD differences between the two states with the hinges around the residues His209, Gly376, and Val380 (Fig.3B). To examine the NBD domain motion in isolation, we ran molecular dynamic simulation with the CHARMM36 force field and TIP3P water (see Methodology) for 10ns and generated 1000 structural snapshots at every 10ps. With these 1000 frames, we produce a phi-psi contour plot on which the Rossmann fold AA residues are superimposed. This shows that most of the residues are rotated in the anti-clockwise direction within the predicted dynamic region (Fig.3C).

Borgnia et al.(2016) described a large movement of His209 during the transition from open to closed state. In presence of GTP in the closed state, His209 swings away from the adenine moiety of NADH (∼10.5Å). In contrast, the distance between His209 and the alpha-phosphate of NADH is ∼4.4Å in the open state, which is comparable with the corresponding distance in the potent activator ADP-bound conformation (Banerjee et al., 2003). In addition to this result, we have identified another critical site near Gly 376 that show key role in the transition (Discussed in the next section).

### GTP-inhibition dynamics involve transition residues and changes in peptide-bond geometry

To determine and characterize the local effects of Gly376 conformational changes, we run the well-tempered metadynamic simulation in PLUMED v2.2.3 (Bonomi et al., 2009), using dihedral angles as collective variables. This simulation predicts the angular and distance distortions in standard peptide-bond geometry for the tripeptide Ala375-Gly376-Gly377 (details about peptide-bond geometry **Supplementary figure 6**). Metadynamics-based free energy surface (FES) calculation for this tripeptide shows three energy picks near *φ* = 0^0^ that were classically described as disallowed regions in the Ramachandran phi-psi plot of protein backbone conformations. In between the picks, two low energy passes are found that parallel to Brereton and Karplus (2015) observations (Fig.4C). The Gly 376 passes the high-energy transition zone (i.e., − 35° ≤ *φ* ≤ + 35°) through this low-energy passes with Ψ ≈ −90°. The Gly 376 apo-conformation (*φ =*47.1°, Ψ = − 109.3°) transits to the conformation (*φ =*− 60.5°, Ψ = − 43.2°) at the closed state via the open state conformation (*φ =*− 53.4°, Ψ = − 43.6°). Along the transition trajectory computed using the molecular mechanics force field CHARMM36, Gly 376 has repeated highest number of time among all the 14 different AA residues trapped in the transition zone, indicating the key role of Gly 376 by the distortion of backbone bond-angles of the tripeptide (**Supplementary table 7**). We have visually checked the conformational occurrences of many transition residues against its electron density map, and found that they are reliably defined (Fig.4A) (See also **Supplementary figure 7**).

**Fig. 6.**
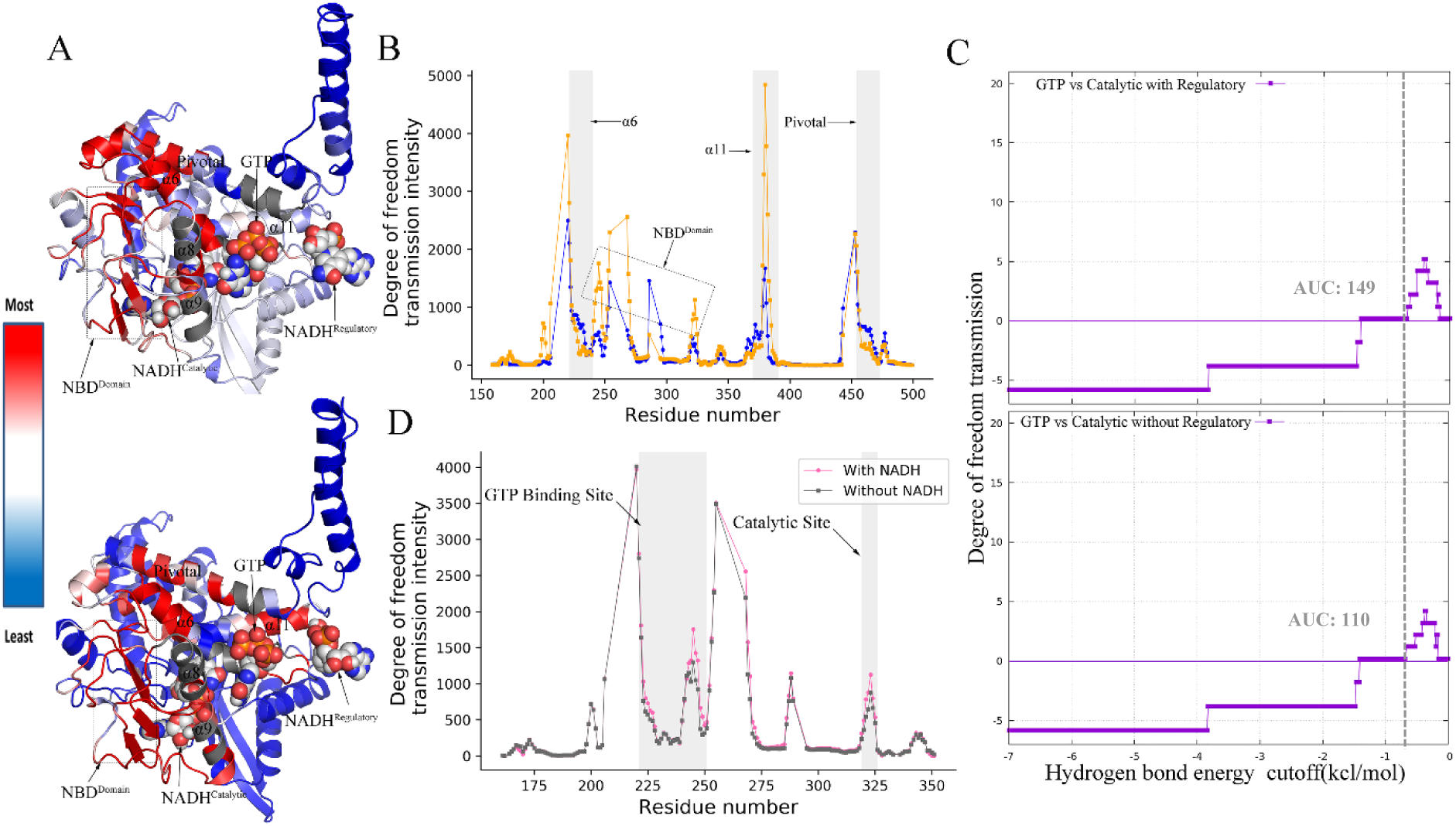
Long range rigidity-based allosteric communications. (A) Perturbing rigidity of pocket defined by atoms within 3.0 Å neighborhood of GTP ligand results in allosteric modification of rigidity and conformational degrees of freedom in the distant catalytic and regulatory sites. Highest degree of freedom transmission intensity (i.e. strongest allosteric effect) is represented by red color and lowest intensity represented by blue. (B) Allosteric hot spots are identified by grey sheds; allosteric intensity at the hot spots is much higher in the closed form (light orange) relative to open conformer (blue). (C) Total area under the allosteric curve (AUC) is higher in presence of regulatory NADH. (D) Synergistic allosteric effects of GTP and regulatory NADH at the catalytic site are significantly higher relative to in absence of regulatory NADH.

**Fig. 7.**
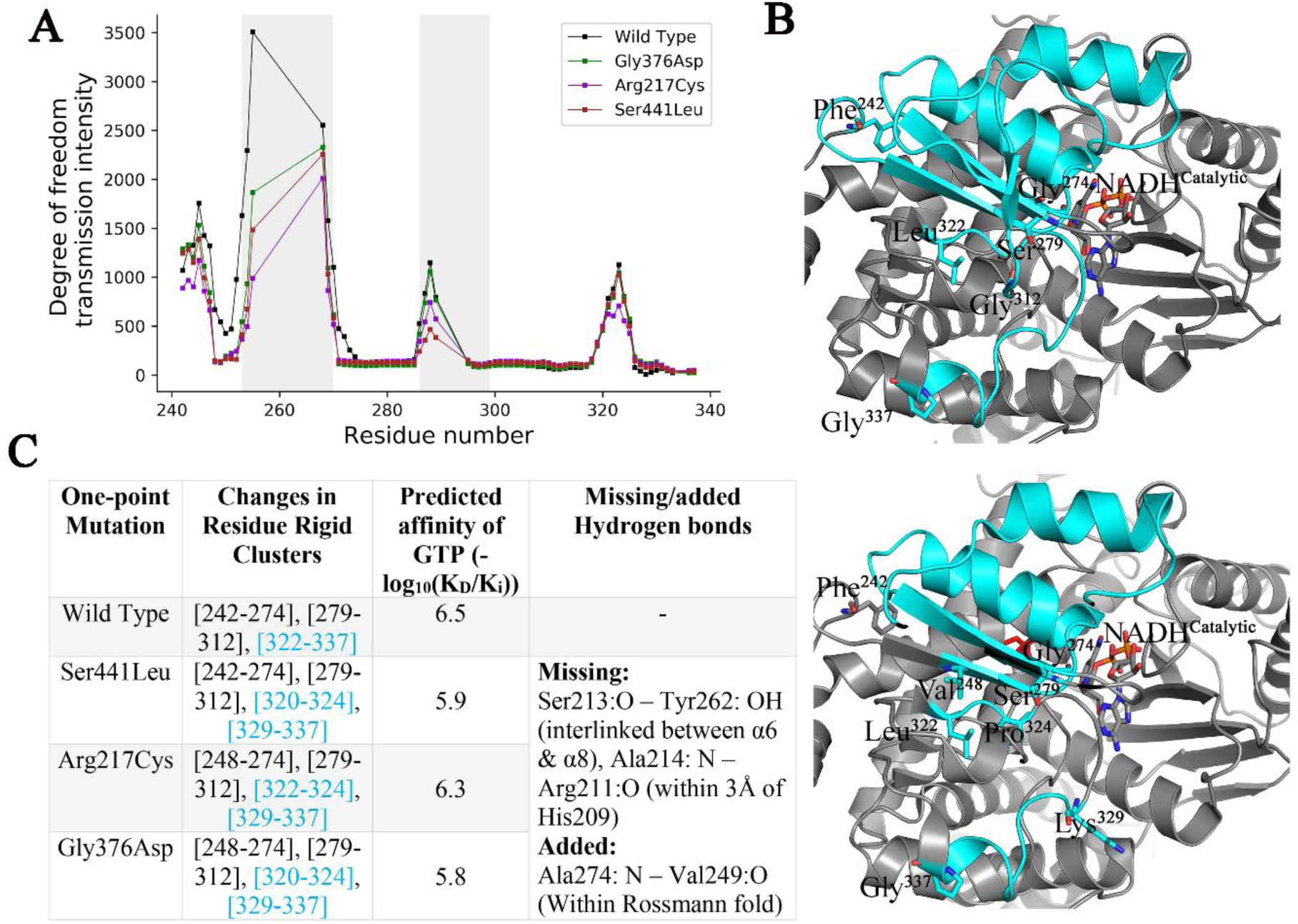
Point-mutations of transient residues, Gly376Asp, show the similar effects with that of the reported mutations Ser441Leu and Arg217Cys in HI/HA syndrome, in which GDH remains insensitive to GTP inhibition. (A) All the stated mutation has undermined the allosteric communications by reducing transmission intensity of the DoFs in-between the rigid allosteric hotspot regions, (B) An allosteric rigid cluster (cyan color) is broken into two parts after the mutations.

Distortions of the distance O^-1^… C and tripeptide backbone bond angles show the generic transition properties and it depends on the torsional angle *φ* along the trajectory. Distortions are *φ*-dependent and can be predicted by a quadratic function of *φ* with the (Fig.4B). Near *φ* =°, the distance is relatively flat and down from the values not in the transition zone. The bond angles ∠C^-1^-N-C _*α*_, ∠O^-1^-C^-1^-N, and ∠C ^-1^-C^-1^-N show systematic variations, with average distortions of roughly ∼2^0^, ∼ 4^0^ and ∼2^0^ respectively from their standard values. The bond angle ∠C^-1^-N-C _*α*_ has the maximum distortion of roughly 6^0^ in the transition zone near to *φ*= 22^0^ (details of other five angle is shown **Supplementary figure 8**) These back-bone bond angles are expressed as a function of *φ* along with cosine function that fit to the data. This smooth curve for both the angles ∠C^-1^-N-C _*α*_ and ∠O^-1^-C^-1^-N display “humped”-type distortions with the hump near to *φ*=0^0^. However, the fitted curve for ∠C_*α*_^-1^-C^-1^-N remains relatively flat in the transition zone −35° ≤ *φ* ≤ + 35°. This analysis therefore shows the key role of Gly376 to cross the energy barriers between the open and closed metastates.

**Fig. 8.**
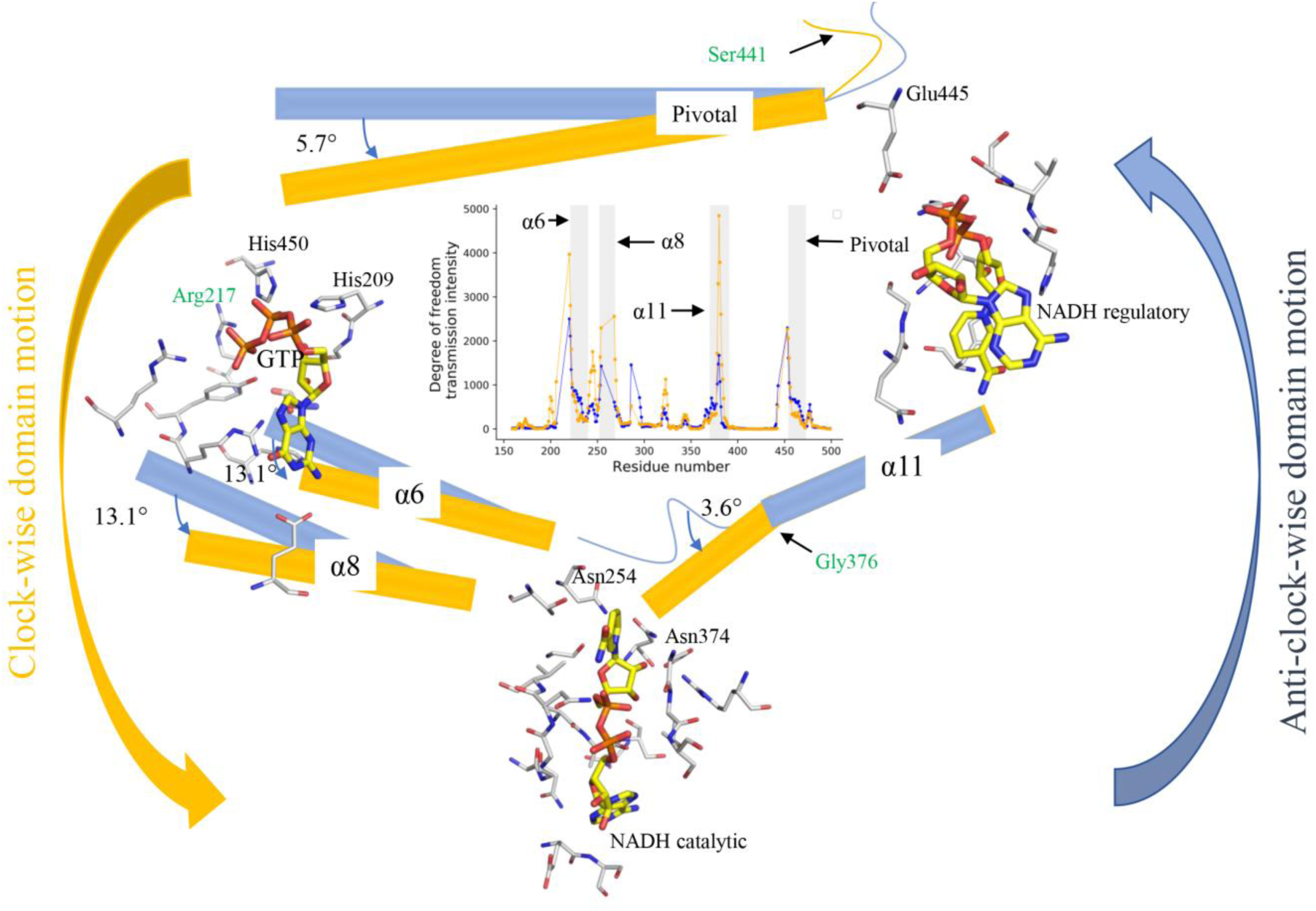
GTP-induced GDH inhibition dynamics: allosterically interacting helices rotate anti-clockwise, in which the transient critical residue Gly 376 has a large conformational change (yellow represents the *closed* and the blue for *open* GDH systems). That generates a torsional force, resulting in tilted extension of the helix *α*11 giving a rotational motion of the Rossmann fold that block the GDH catalytic cleft. The dominant HI/HA mutations Ser441Leu and Arg217Cys break this allosteric network, causing GTP-insensitivity and thereby over-activity of GDH. GDH_Gly376Asp_ mutant shows similar effect, inferring the key role of Gly 376 for the inhibition of GDH activity by GTP.

### GTP-binding forms an allosteric nexus

GTP-binding triggers an allosteric response connecting the three distant regions of GTP binding, NADH regulatory, and catalytic sites and thereby forming an allosteric nexus (Fig.6A). To determine the associated allosteric network we use graph theoretical method FIRST (Jacobs et al., 2001) which is based on pebble game algorithm and techniques in rigidity theory. FIRST efficiently decomposes the protein into rigid clusters and connecting flexible regions. Activation and functioning of allosteric network are indicated by the formation of a largest rigid cluster (LRC) that encapsulates the AA residues from a small neighborhood of these three distant sites (Fig.5). Rigid cluster decomposition of apo, open and closed GDH structures with the acceptable H-bond energy cutoff of -0.5 kcal/mol (**Supplementary figure 5**) shows that GDH closed conformer has the largest LRC, roughly about ∼1.8 times bigger than the LRC in the open form, with lowest total independent degree of freedoms (DOF). As expected GTP-bound open conformer has the smallest LRC and highest DOFs among all the three structures, referring to formation and activation of allosteric network through conformational rearrangements of LRC residues on the onset of GTP binding. Moreover, we note that the LRC of GDH closed conformer includes approximately 2-fold more helical structures than its open form, making it much less flexible and potential for intense allosteric communication. Consequently, total number of rigid clusters has significantly reduced when it transformed from open to closed form. In particular, α11 that play important role in the GTP-inhibition dynamics has contributed 84% for LRC formation in the closed form in contrast to its 33% inclusion in the open form (**Supplementary table 6**).

Binding of GTP imposes local constrains by introducing two strong H-bonds with Arg 261. Furthermore, this local constraint has become more stringent in presence of NADH cofactor at the regulatory site which includes the formation of two more H-bonds with Tyr 262 and His 209. This GTP-induced perturbation in the local conformational rigidity within the 3.0 Å neighbourhood of GTP is transferred to other remote sites allosterically, resulting in a large NBD motion that closes catalytic cleft. To examine this allosteric effect quantitatively, we apply the rigidity transmission allostery (RTA) algorithm (Kim et al., 2017) that predicts and quantifies the potential allosteric communications between the GTP binding site and the other remote sites. Residues were labeled as allosteric hot spots based on the intensity of transmission of degrees of freedom (DOF). Initially, both the GTP-bound GDH open and closed conformers were decomposed into rigid clusters and flexible regions using the program Floppy Inclusion and Rigid Substructure Topography (FIRST) and then RTA algorithm was applied to calculate if DOF propagate through protein structure as a result of rigidifying the GTP binding pocket. The RTA algorithm evaluates if perturbation of rigidity in GTP-binding neighborhood has caused changes in rigidity and conformational degrees of freedom in the catalytic and regulatory sites (i.e. allosteric transmission). Overall, RTA analysis shows that average intensity of transmission of DOFs is significantly higher in the closed state compared to open GDH conformer (Fig.6B). Excluding the antenna domain that holds a large intrinsic motion due to its long protruding geometry and slight conformational changes in its immediate vicinity, residues 379-385 residing on the helix α11 are displayed to be allosteric hot spots, indicting the transfer of local rigidity to regulatory NADH sites. Simultaneously, residues 220-223 resided on α6 are allosteric hot spots, including the residue Asn 254 that formed three H-bonds with the catalytic NADH. RTA analysis therefore shows allosteric communication between the binding sites of GTP, catalytic NADH, and regulatory NADH. This triangular allostery triggered by GTP binding synergistically affects the GDH catalytic activity and eventually blocks the catalytic cleft.

To probe the allosteric role of regulatory NADH, we further applied the RTA algorithm on the GTP-bound GDH system in presence and absence of regulatory NADH. Perturbation effects of rigidity within 3.0 Å neighbourhood of GTP on the catalytic sites are calculated based on allosteric intensity for both cases and then area under the allosteric (degree of freedom transmission) curve (AUCs) (Jeliazkov et al., 2018) are calculated numerically that indicate the overall strength of allosteric intensity in presence and absence of regulatory NADH (Fig.6C,D). AUC in presence of regulatory NADH is significantly higher than the AUC in absence of regulatory NADH, implying the specific role of regulatory NADH as positive allosteric regulator. Moreover, synergistic effects of GTP and NADH are reflected in the DOFs transmission intensity particularly at catalytic region which are much higher compared to the DOF intensity transmitted without regulatory NADH.

### Mutation of allosteric hot spot residues alter the allosteric communications

In order to examine how the triangular allosteric network responds to the mutations of key residues, we prepared three GDH mutants *in silico* using Chimera rotamers tools (Pettersen et al., 2004): GDH_Ser441Leu_, GDH_Arg217Cys_, and GDH_Gly376Asp_. As reported by Stanley (2011), GDH_Ser441Leu_ mutant is associated with the HI/HA syndrome and is the result of a substitution from Serine TCG to Leucine TTG codon in the *GLUD1* gene that encode the enzyme GDH. Recent study further noted that Ser441Leu is the most frequent sporadic mutation among the HI/HA patients (Grimaldi et al., 2017) (**Supplementary table 5**). This mutant shows over-activity of GDH caused by extreme insensitivity to GTP inhibition; however, the molecular explanations of this insensitivity remain unknown. To probe the effects of mutants on allosteric network, we repeated the RTA allosteric analysis of the GDH mutants. The RTA calculations with Ser441Leu mutation has undermined the allosteric communications by reducing transmission intensity of the DoFs in-between the rigid allosteric hotspot regions (Fig.7A). Furthermore, mutation Ser441Leu destabilized rigid cluster [resi.322-resi.337] in the closed conformation where rigid cluster broke off into two parts. In particular, this caused a loss of two H-bonds between Ser213.O-Tyr262.OH and Ala214.N-Arg211.O (Fig.7B,C). This change in the rigidity disconnects the catalytic region from the GTP-binding sites, resulting in reduced motion of the Rossmann fold that fails to block the catalytic cleft as it observed in the closed type GDH conformation.

In parallel to Ser441Leu mutation located in the vicinity of the GDH antenna, GDH_Arg217Cys_ mutation is also associated with HI/HA and leads to insensitivity to GTP inhibition (Stanley, 2011). Although Arg 217 is located in the GTP-binding domain, GDH_Arg217Cys_ mutant shows similar effects on the allosteric network, including the inclusion and exclusion of the same H-bonds (**Supplementary table 8**). Notably, binding affinity of GTP remains unaltered in both the mutations. As the residue Gly 376 -- a key transient residue responsible for the transition from open to closed conformation, Gly376 is substituted by Asp following the list of reported HI/HA associated mutations of GLY. Interestingly, it is noted that the mutant GDH_Gly376Asp_ mimics the similar effects on the allosteric network. Moreover, we performed the metadynamics on the GDH_Gly376Asp_ mutant and then recalculated the free energy surface (FES) for the tripeptide Ala375-Asp376-Gly377. The predictions show the loss of the two low energy passes as noted for the Gly376, which has prevented the crossing of the energy barriers by Asp 376 (Supplementary figure 8D). Therefore, this *in silico* mutational analysis reinforces the key role of the transient residue Gly 376 for maintaining the GTP-induced inhibition dynamics.

## Conclusion

The GDH regulatory systems involve a variety of protein effectors and many small allosteric modulators that build up a complex allosteric network. Recent discovery of HI/HA syndrome show GDH dysregulation through insensitivity to its usual inhibitor GTP and thereby over-activity of GDH (Stanley 2011; Grimaldi et al., 2017). Here we have illustrated an allosteric regulation of GDH by GTP and provide a molecular explanation for the dominant HI/HA mutations Ser441Leu and Arg217Cys that causes GTP-insensitivity. The details of regulatory insights of GDH system are also important to understand the fundamental link between carbon and nitrogen metabolism, as the GDH catalyzes the deamination reactions that supplies the alpha-ketoglutarate as inputs to TCA cycle. In this study we have shown how a purine nucleoside triphosphate, Guanosine triphosphate (GTP), binding establishes rigidity-based allosteric communication with the regulatory coenzyme NADH and then synergistically inhibits the oxidative deamination activity of mammalian GDH by stabilizing its closed conformation. This work, therefore, contributes to better understanding of the complex link between protein flexibility, allostery and protein function.

In particular, we show that GTP-binding triggers interhelical interactions and subsequent anticlockwise motion of nucleotide-binding domain (NBD) involving the Rossmann folds that eventually close the catalytic cleft. This NBD motion is controlled through a triangular allosteric network linking the GTP binding, regulatory NADH, and catalytic sites. It is further noted that the residue Gly 376, a critical residue located at the N-terminal of the alpha-helix 11, play an important role through local geometric changes in the tripeptide bond. These alterations result in a tilted extension of the helix 11 towards the NBD and generated a torsional force that provides the anti-clockwise motion to the Rossmann fold. *In Silico* mutational comparative analysis shows the similar allosteric effects of Gly376Asp mutation, as observed for the dominant HI/HA mutations. Moreover, the analysis illustrate the role of the cofactor NADH as a positive allosteric modulator (PAM), increasing the allosteric effect by increase in intensity of transmission of the degree of freedoms (DOFs) through the allosteric network. In addition, regulatory NADH enhances GTP binding affinity, with synergistic inhibitory effects on the GDH catalytic domain.

## Methods

### Systems construction for MD-Simulation

The open form of NADH- and GTP boundbound GDH system (PDB id: 3jd3) cryo-EM structure was chosen and ligands, water molecules, other solvent and unwanted heterogens were solved and removed from structure by PDBFixer at pH 6.8 (Eastman et al., 2017). The fixing structure was collected without adding missing hydrogen atoms. All hydrogen atoms were placed with the GROMACS tool pdb2gmx. All three separate ligands GTP and both regulatory and catalytic NADH were placed into Avogadro program (Hanwell et al., 2012) for adding hydrogen and producing .mol2 file. We have used standard sort_mol2_bonds.pl perl program code for fixing the bond listing and other small error like those atoms were assigned different residue name and number. Topology and parameter files were constructed from final corrected structure with the help of CGenFF web server (Vanommeslaeghe et al., 2010). Finally, Gromacs formatted topology was constructed using the cgenff_charmm2gmx.py script with CHARMM36 force field (Huang et al., 2013) and added with protein structure topology file to generate protein-ligand complex.

### MD simulation and parameters

Molecular Dynamics simulations were organized and carried out with GROMACS v5.1.4 (Abraham et al., 2015) simulation package of version 5.1.4. Proein-ligand complex was solvated in a rhombic dodecahedral unit cell consisting of 41638 water molecules, 90 Na^+^ and 82 Cl^-^. The complex was modelled with CHARMM36 force field and TIP3P water. All the MD simulation steps include energy minimization, equilibrations and production were performed by 2fs time steps. The system was subjected to maximum 50000 steps of steepest decent minimization or up to the maximum force dropped less than 1000 KJ/mol/nm to remove unfavorable energy contacts. Equilibration were carried out by two successive blocks of simulation with 0.1 ns each in the NVT and NPT statistical mechanical assembles where all bonds including heavy atoms-h bonds were constructed. The production simulation was performed with randomized initial velocities of all atoms total up to 10ns at a temperature of 300K. These simulation was carried out in the NPT assembles by heating to 300K with the Nosé-Hoover thermostat (Hoover, W. G, 1985). Pressure (1 bar) was maintained using the Parrinello-Rahman barostat (Parrinello et al., 1981) with coupling constant of 2ps. Coulomb long-range electrostatics interactions were evaluated using the smooth particle-mesh Ewald method(Darden et al., 1993) with cubic interpolation of order 4 and a Fourier grid spacing of 0.16 nm. Bonds to hydrogen atoms were constrained with LINCS (Hess et al., 1997).

### Metadynamics

Well-tempered metadynamics simulation (Barducci et al., 2008) was performed on Gly376 by GROMACS patched with PLUMED v2.2.3 (Bonomi et al., 2009), using as collective variable (CV) the backbone dihedral angle phi and psi. The stride of Gaussian deposition was updated over every 500 time steps while the height of the Gaussian potentials was set in 1.2 kJoule/mol. The width of the Gaussian potentials for each CV was specified to 0.35 rad at temperature 300K.

### Curating the MD trajectory data for High-energy pass and protein geometry

From the MD trajectory data, 1000 structural snapshots were created over every 10ps time step. On the basis of these snapshots, all the observation having their φ torsion angles in the high energy pass region (−35 < φ < +35), were manually curated. Reliability of the structural conformation was tested on the basis of visual assessment of the fit to their Electron Microscopy (EM) map with Chimera v1.3 package Fit in Map (Pettersen et al., 2004). Reliable structures were separated whose φ torsion angle belongs to high energy pass region. Unreliable or close to unreliable residues were also included in this range to present of alternative conformations. All this curated observed residues in this range extracted from MD trajectory data by writing a custom script in GROMACS and GNU plot. Some specific geometric details were also calculated for all residues from each protein snapshot, excluding those residues not having two residues of both sides of it. The geometrical quantities include backbone torsion angles, bond angle and the O^-1^…C distance.

### GNM and ANM analyses for protein dynamics

Anisotropic network model (ANM) (Eyal et al., 2015) and Gaussian network model (GNM) (Li et al., 2016) analysis was done using DynOmics web server (http://dynomics.pitt.edu/) (Li et al., 2017) and Bio3D version 2.3.0. of R package (Grant et al., 2006). Relative mode of motion was identified along the principle mode using GNM. Block of residues were divided into separated domains based on the direction (+/-) of their moment in the slowest modes. The sign (+/-) of the elements (residues) based on the selected mode eigenvector. Same sign indicates the structural regions that have correlated (positive) motion and opposite sign linked with anticorrelated (negative) motion. The theory of ANM and GNM can be found in ref. (Eyal et al., 2015) and ref. (Li et al., 2016), respectively. Overlap analysis was carried out using Bio3D to identify which modes contribute a given conformational change. Structure visualition, analysis and animation were performed in the NMWiz (Normal Mode Wizard, Version 1.0) tools of VMD (Bakan et al., 2011).

### Rigidity based Allosteric Computations

The three way allosteric transmission from NADH regulatory site to catalytic site through GTP binding site, is observed using RTA algorithm, a computational approach based on rigidity theory (Whiteley et al., 2005) and an extension of FIRST method (Jacobs et al., 2001). The RTA algorithm was first initiated in 2012 (Sljoka et al., 2012) and further developed and discussed by Whiteley et al. (Finbow-Singh et al., 2013). This algorithm extends the pebble game algorithm to predict whether local perturbation of rigidity at one site of the protein, transmit across the structure to change the rigidity of the second distant site. FIRST generates a constraints network from the coordinates of the protein structure in terms of nodes (atoms) and edges (covalent bonds, hydrogen bonds and hydrophobic interactions). Hydrogen bonds were ranked according to their energy strength using Mayo potential (Jacobs et al., 2001), whereas the value of energy strength was selected in such way that the bonds strength below this cut off were ignored. First then applies the rigorous mathematical theory of rigid and flexible molecular structure and pebble game algorithm (Jacobs et al., 1997) calculates the degree of freedom of motion to rapidly decompose a protein into rigid clusters and flexible region. We applied RTA analysis determines whether perturbation of rigidity and conformational degrees of freedom at NADH regulatory site could propagate through the protein structure (Pivotal helix, α11 and Rosenman fold) inducing the quantifiable changes in rigidity and available number of degree of freedom at a second distant site (GTP binding region), hence result in allosteric transmission. The number of conformational degree of freedom at the GTP binding region was calculated before and after a sequential perturbation of rigidity of all residue at the regulatory NADH binding region. To understand the role of regulatory NADH, the changes of number of conformation degree of freedom also measured after removing the regulatory NADH from the structure.

### Area Under Curve (AUC)

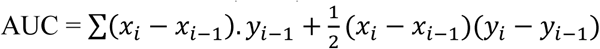

The AUC were approximated by simple integral (alike to trapezoidal integration) where the first term defines a rectangle and second term denies a tringle. (Jeliazkov et al., 2018).

### Rigidity-based transmission allostery

Missing hydrogen atom were added to Cryo-EM structure using WHAT IF web server (http://swift.cmbi.ru.nl/servers/html/htopo.html) for rigidity and allostery analysis. We sequentially perturbed the rigidity of a window of the three-consecutive residue (r, r+1, r+2) starting from N-terminal for calculating degree of freedom that could be transmitted to the GTP binding region. Perturbation of rigidity means insertion of additional constraints (edges, removal of degree of freedom) to rigidify it. Transmission of degree of freedom indicates a subsequent change of degree of freedom at GTP binding region, consequences the presence of rigidity-based allostery. We define binding site as the residues belongs to 3Å neighboured of GTP and NADH. The RTA algorithm computes the tansmission of degree of freedom between two sites where as NADH regulatory site as a site A and GTP binding site as site B. We calculate the number of available conformation degree of freedom at site A, B and the union of sites A and B. This calculation was repeated by successively omitting week energy constraints upon increments of 0.01kcl/mol. We count the degree of freedom at given energy cut-off as A_max_, B_max_ and AB_max_ respectively (using pebble game algorithm). Finally, The maximum number of degree of freedom could be transmitted from A to B (denoted by DOF_AB) were also calculated by obtaining DOF_AB = A_max_ + B_max_ - AB_max_ – 6 (the trivial degree of freedom corresponds to rigid body motion is 6). Positive DOF_AB indicates the sites A and B are involved in rigidity based allosteric transmission and the maximum number of degree of freedom that can be transmitted from A to B is DOF_AB. Allosteric transmission for some residue r refers a transmission curve generated from the average DOF_AB for three consecutive windows containing r [i.e. (r-2, r-1, r), (r-1, r, r+1) and (r, r+1, r+2)] as a function of energy cut-off. To observe the intensity of allosteric transmission for each window we computed the area of under the transmission curve. Therefore the intensity of allosteric transmission for a residue r was calculated from the average intensity of three consecutive windows containing the residue r. Therefor transmission of allosteric intensity considered the no degree of freedom that could be transmitted and persistence of the transmission as a function of energy strength. Freedom. Moreover, allosteric transmission persists of a wide range of energy cut-off indicates robust allosteric communication that means small changes in hydrogen bonding network will not significantly affect the allosteric transmission. (Kim et al., 2017; Ye et al., 2018).

### Mutational effect on allostery

Protein structure was prepared for point mutation using Chimera rotamers tool (In structure editing section) (Pettersen et al., 2004; Rashid et al., 2018). Upon mutation, residue conformation was chosen according to highest probability from the rotamer library (probability is taken from the library that was not affect the structural environment but changed in Phi and Psi angle when Dunbrack library was used). We have chosen two mutations (Ser441Leu and Arg217Cys) from different hotspot position (regulatory NADH and GTP binding site) with highest frequency of HI/HA mutations according to Stanley et al. (Stanley CA., 2011). Then we mutated Gly376 with possible replacement having seen in HI/HA syndrome patent (Arg, Cys, Ser, Asp, Val) and we were chosen Asp for mimicking the mutational effect on structural rigidity with Ser441Leu and Arg217Cys. After mutation, each structure was minimised for removing classes and contact up to 1000 steps using Steepest decent method with step size 0.02Å and none of the atom was fixed. After adding hydrogen, we also added charges using Amber ff99SB force field. We also defined ligand charges using ANTICHEMBER (Wang et al., 2006) for minimisation. All this process was completed using chimera Minimize Structure tool. Before used this mutated structures for further analysis, we removed all the hydrogen from the structure to maintain the method of adding polar hydrogen. To understand the effect of mutation on allosteric transmission, transmission of allosteric intensity was applied with RTA and changes in rigid clusters using FIRST (compared with wild type with mutated structures using FIRST at hydrogen bond energy strength -1.0Kcl/mol), we carried out same process (as described above) on all mutated structures (Gly376Asp, Ser441Leu and Arg217Cys). Well-tempered metadynamics simulation (Barducci et al., 2008) was performed up to 10ns on Gly376Asp by GROMACS patched with PLUMED v2.2.3 (Bonomi et al., 2009) to understand the high energy pass and protein geometry upon mutation (Details in Supplementary).

## Acknowledgments

Our thanks to project students Ayushi Bhutra, Govind Swami, Neha Bhookar, and Kartik Sharma of the School of Mathematics, Statistics and Computational Sciences, Central University of Rajasthan (CURAJ) for their support while completing a part of computational study of this work. This research was supported by SERB-EMR grant (file no. EMR/2015/001671) and by the Central University of Rajasthan; M.R. received Ph.D. fellowship from the Central University of Rajasthan and S.B. received Ph.D. fellowship from CSIR-HRDG.

## Author Contributions

S.B., M.R., A.S., A.C. designed the study and conducted computational experiments and analytical analysis. A.C., G.-Q.S., A.B.M., A.S., B.L. have reviewed and written the article.

