## Supplementary file for "Allosteric regulation of Glutamate dehydrogenase deamination activity"

**[2]Department of Mathematics, North University of China, Shanxi, China.**

**[3]Institute of Theoretical and Experimental Biophysics, Pushchino, Russia.**

**[4]University of California, Riverside, USA.**

**[5]Institute for Theoretical Biology, Humboldt University, Berlin, Germany.**

**[6]RIKEN Center for Advanced Intelligence Project, Tokyo, Japan.**

**[7]Department of Chemistry, University of Toronto, Canada.**

***Corresponding author:**
Amit Chakraborty


**Lead contact:**
Amit Chakraborty


| **Helix No** | **Apo (Helix size)** | | | **NADH+GTP+GDH open (Helix size)** | | | **NADH+GTP+GDH closed (Helix size)** | | | **Correlation Coefficient(r)** |
|---|---|---|---|---|---|---|---|---|---|---|
| | Residue Id | Length (Å) | Radius (Å) | Residue Id | Length (Å) | Radius (Å) | Residue Id | Length (Å) | Radius (Å) | |
| α1 | 8-29 | 33 | 2 | 8-32 | 35.5 | 2.4 | 8-30 | 34.1 | 2.3 | 0.79577 |
| α2 | 38-53 | 25.2 | 1.8 | 39-53 | 23.5 | 1.9 | 37-53 | 25.7 | 1.9 | |
| α3 | 100-115 | 23.6 | 1.9 | 100-118 | 28.1 | 1.9 | 100-118 | 27.6 | 1.9 | |
| α4 | 139-154 | 24.9 | 1.9 | 139-155 | 26.1 | 1.8 | 139-155 | 26.1 | 1.8 | |
| α5 | 172-184 | 19.7 | 1.9 | 172-184 | 19.6 | 1.9 | 172-187 | 23.2 | 2.0 | |
| α6 (intermediate of pivotal, α8 and α9 helix) | 213-237 | 32.6 | 2.7 | 213-224 | 19.2 | 1.8 | 213-237 | 32.2 | 2.7 | 0.25427 |
| α7 | -------- |  |  | 229-234 | 8.1 | 2.0 | ----------- |  |  |  |
| α8 | 253-265 | 18.7 | 1.9 | 255-265 | 17.3 | 1.8 | 253-265 | 19.1 | 1.9 | |
| α9 | 287-299 | 16.9 | 2.1 | 287-298 | 16.4 | 2.0 | 287-298 | 16.1 | 2.0 | |
| α10 | 353-361 | 14.2 | 1.8 | 353-363 | 16.1 | 1.8 | 353-363 | 15.9 | 1.9 | |
| α11 (elongated helix) | 376-391 | 23.5 | 1.9 | 369-388 | 31.5 | 2.0 | 369-388 | 31.2 | 2.0 | |
| α12 (Antenna helix) | 398-425 | 38.7 | 2.4 | 398-407 | 15.9 | 1.8 | 398-425 | 42.2 | 2.3 | 0.21784 |
| α13 (Antenna helix) | --------- |  |  | 409-421 | 19.3 | 1.8 | ---------- |  |  |  |
| α14 (Antenna helix) | 434-442 | 13.1 | 1.9 | 433-442 | 13.6 | 2.0 | 433-439 | 10 | 1.9 | |
| α15 (pivot helix) | 444-471 | 42.1 | 1.9 | 444-471 | 42.2 | 1.9 | 444-471 | 41.1 | 1.9 | 0.79394 |
| α16 | 476-497 | 33.3 | 1.9 | 476-497 | 33.2 | 1.9 | 476-496 | 33.1 | 1.9 | 0.03860 |

**Supplementary table 1: Detail information of helices.** Table represents helix number, helix length, radius and corresponding correlation coefficient (r) compare with the open structure. Helix number is introduced from Stride Services (Heinig et al.,2004). Helix length and radius is calculated from Chimera software. Low deviation correlation coefficient indicates the lack of similar pattern between helices (open to closed). Table indicates α1- α5 and α15 have high level of correlation compare to helices α6 to α11 and Antenna helices. In contrast, the table suggest a lack of similarity in NBD (α6 – α11) and Antenna region indicates helical rotation and translation.

**Notes: Deviation of Cα atoms:**

Cα deviation movement is measured using formula:

$$\sqrt{\frac{1}{n}\sum_{1}^{n}((x_i - x_0)^2 + (y_i - y_0)^2 + (z_i - z_0)^2)}\,,$$

where $(x_i, y_i, z_i)$ are the position of the Cα atoms of every residue within GDH (3jd3 open and 3jd4 closed). The deviation is measured in all cases from unbound GDH structure (3jcz). Deviation is compaired and visualize with different conformation of GDH using small code of Python. Helix number also included with this comparison to understand the region of changes in deformation of Cα atom between different conformations of GDH.

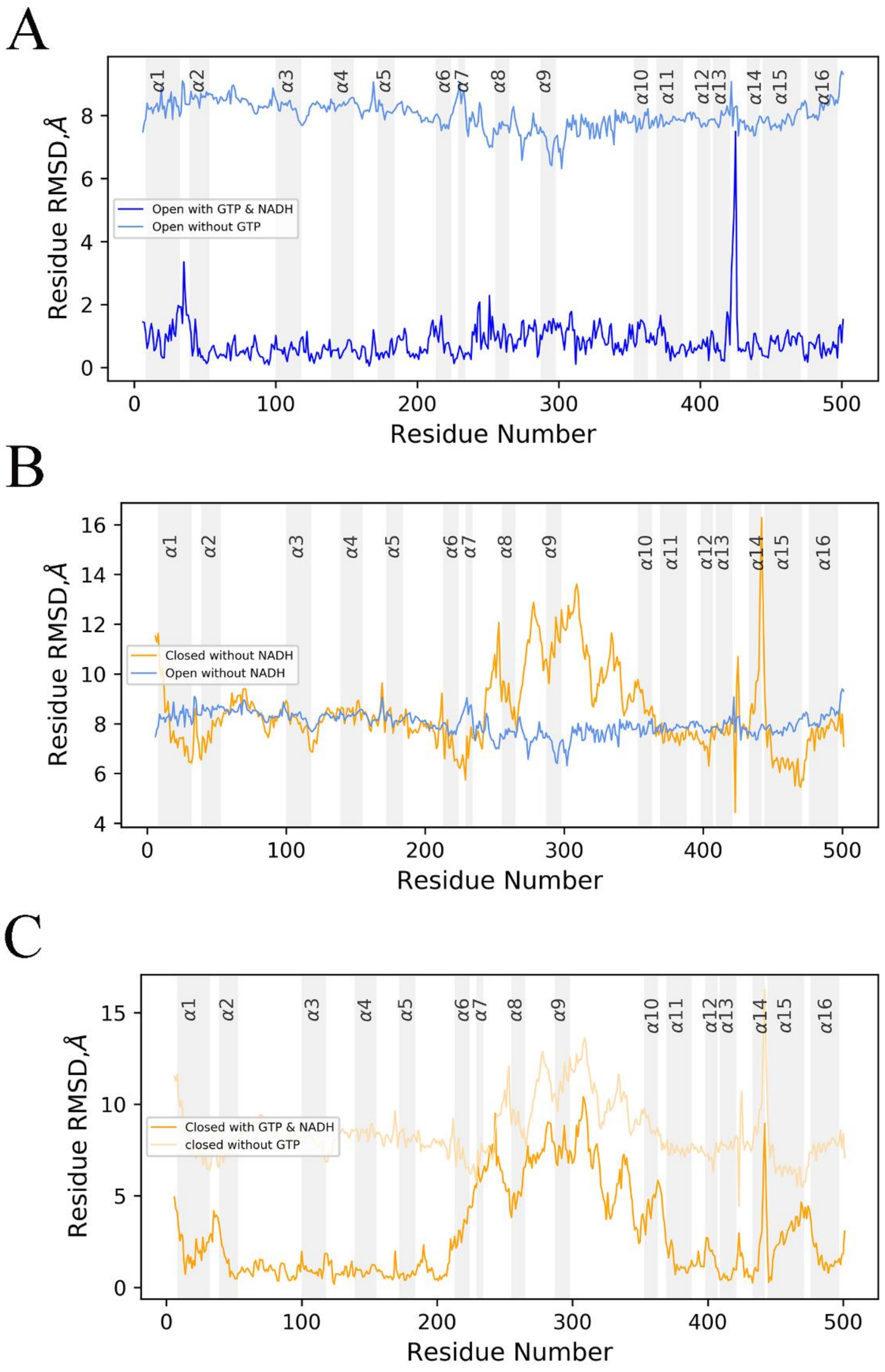


**Supplementary figure 1: Compare of Cα deviation**. Figure showing the individual Cα deviation of different structures (GDH with NADH closed, GDH with NADH open, GDH with NADH and GTP open, GDH with NADH and GTP closed) superimposing with Apo structure (GDH only). (A) Plot indicates the effects of GTP on open structures (B) Figure also indicates the effects of GTP on open as well as closed structure (GDH with NADH open and GDH with NADH closed). (C) Plot compares individual residue deviation between closed structures with and without GTP. Helices are indicated with shaded region and all the figure is generated using Python code.

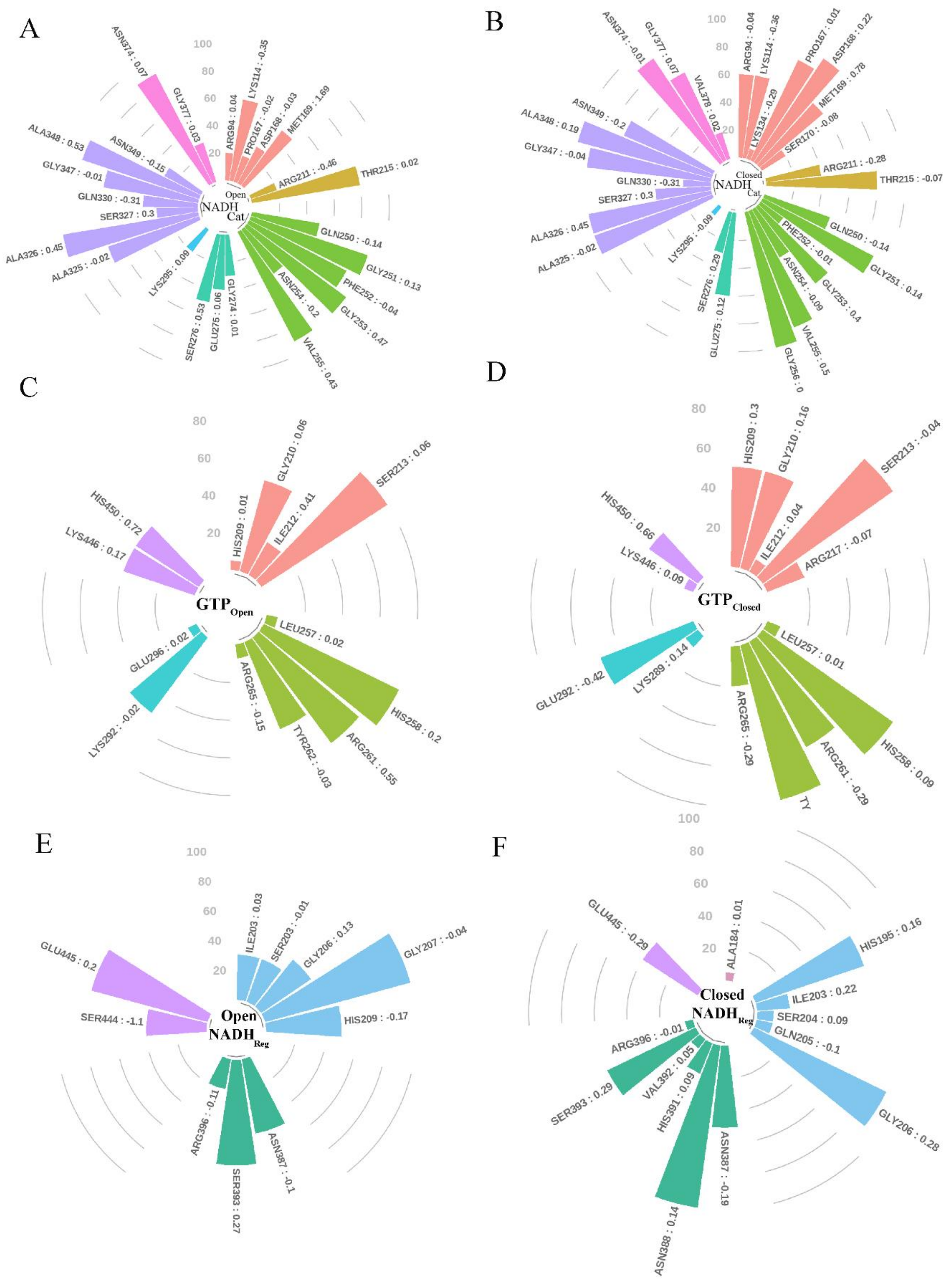


**Supplementary figure 2: Interaction between cofactors NADH, inhibitor GTP with GDH.** Plot showing detail information of interfaces residues with ligands at two different conformations (open and closed). Group wise colours designates with alpha helices or outside of alpha helices. Each bar represents the contribution of buried surface area (BSA) of interfacing residues with ligands and fraction numbers at top of the head of each bar showing solvation energy effect ($\Delta^{i}G$ Kcl/mol). (A), (B) figure showing the interaction between interface residue of protein and catalytic NADH at open and closed confirmation. Number of interfacing residues (29) in closed conformation much higher than open (26) at catalytic site. Medium purple colour bars indicate the residue in NBD domain but not in alpha helices however violate and lime green bars situated at α11 and α8 respectively. (C), (D) represents the interfacing residues of protein with GTP. Percentage of buried surface area of His 209, Arg261 and Tyr 262 are sufficient increased in closed structure compare to open indicate

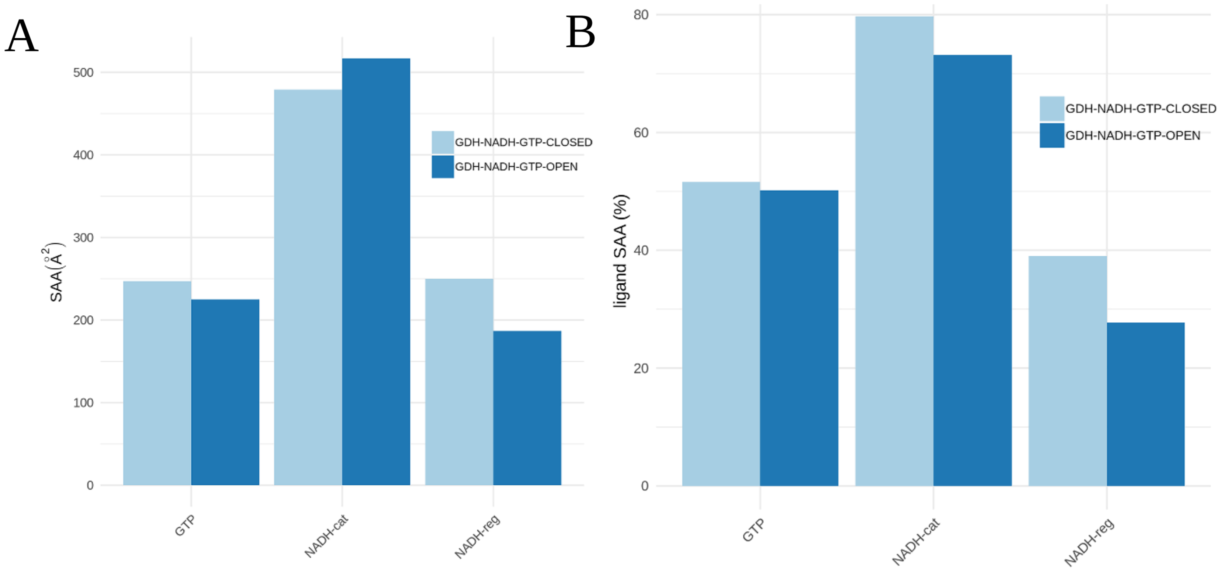


**Supplementary figure 3: Solvent Accessible Area (SAA) of protein and ligand in-between the protein ligand interfaces**. (A) figure represents Solvent Accessible Area (SAA) of protein interfaces between GTP, NADH catalytic and NADH regulatory site however figure (B) showing the SAA (%) of GTP, NADH catalytic and NADH regulatory between protein. The solvent accessibility of protein at NADH catalytic site is significantly higher for open structures indicates the flexibility of protein at catalytic site compare to closed one.

**(A) 3jd3 open structure: Interface**

| Structure1 | | | | Structure2 | | | | | | |
|---|---|---|---|---|---|---|---|---|---|---|
| **Range** | $N_{at}$ | $N_{res}$ | Surface ($Å^2$) | Range | Nat | Nres | Surface ($Å^2$) | Interface Area($Å^2$) | ΔG Kcl/mol | $N_{HB}$ |
| **Chain A** | 73 | 28 | 23318 | NAI: Catalytic | 42 | 1 | 825 | 560.0 | -7.8 | 8 |
| **Chain C** | 30 | 10 | 23318 | NAI: Regulatory | 20 | 1 | 773 | 200.8 | -1.9 | 2 |
| **Chain A** | 34 | 13 | 23318 | GTP | 24 | 1 | 540 | 248.0 | -5.4 | 3 |

**(A) 3jd4 closed structure: Interface**

| Structure1 | | | | Structure2 | | | | | | |
|---|---|---|---|---|---|---|---|---|---|---|
| **Range** | $N_{at}$ | $N_{res}$ | Surface ($Å^2$) | Range | Nat | Nres | Surface ($Å^2$) | Interface Area($Å^2$) | ΔG Kcl/mol | $N_{HB}$ |
| **Chain A** | 80 | 31 | 22631 | NAI: Catalytic | 43 | 1 | 802 | 559.2 | -6.4 | 8 |
| **Chain A** | 40 | 14 | 22631 | NAI: Regulatory | 28 | 1 | 829 | 286.9 | -3.4 | 2 |
| **Chain A** | 34 | 14 | 22631 | GTP | 26 | 1 | 573 | 271.4 | -1.4 | 7 |

**Supplementary table 2: Comparison between protein and ligand interfaces.** The above table Structure1 represent GDH and Structure2 represent cofactors NADH and inhibitor GTP. $N_{at}$, $N_{res}$ indicate number of atoms and number of residues involved in interface area between protein and cofactors as well as inhibitor. $N_{HB}$ indicates no of hydrogen bonds present at interfaces and ΔG (Kcl/mol) is solvation energy for folding. All information were extracted from PDBePISA (https://www.ebi.ac.uk/pdbe/pisa/).

A **Catalytic NADH interface: Hydrogen bonds**

| *No* | Structure1 | Dist (Å) | Structure2 |
|---|---|---|---|
| *1* | A: ASN 254 [N] | 2.92 | A: NAI 602 [O2A] |
| *2* | A: PHE 252 [N] | 2.88 | A: NAI 602 [O3B] |
| *3* | A: GLY 253 [N] | 3.11 | A: NAI 602 [O3B] |
| *4* | A: SER 276 [N] | 2.47 | A: NAI 602 [O2B] |
| *5* | A: ASN 254 [N] | 2.76 | A: NAI 602 [O3] |
| *6* | A: ASN 254 [ND2] | 3.03 | A: NAI 602 [O2N] |
| *7* | A: THR 215 [OG1] | 3.89 | A: NAI 602 [O7N] |
| *8* | A: GLN 250 [O] | 3.61 | A: NAI 602 [N3A] |

**Regulatory NADH interface: Hydrogen bonds**

| *No* | Structure1 | Dist (Å) | Structure2 |
|---|---|---|---|
| *1* | A: SER 444[OG] | 3.40 | C: NAI 601[ O3D] |
| *2* | A: HIS 209[ NE2] | 3.84 | C: NAI 601[ O2D] |

B **Catalytic NADH interface: Hydrogen bonds**

| *No* | Structure1 | Dist (Å) | Structure2 |
|---|---|---|---|
| *1* | A: SER 170 [N] | 2.92 | A: NAI 601 [O1A] |
| *2* | A: ASN 254 [N] | 3.26 | A: NAI 601 [O2A] |
| *3* | A: ASN 254 [N] | 3.68 | A: NAI 601 [O3] |
| *4* | A: ASN 254 [ND2] | 3.54 | A: NAI 601 [O2N] |
| *5* | A: ARG 94 [NH2] | 3.85 | A: NAI 601 [O3D] |
| *6* | A: ASN 349 [N] | 3.78 | A: NAI 601 [O2D] |
| *7* | A: ARG 94 [NH2] | 3.03 | A: NAI 601 [O2D] |
| *8* | A: ASN 349 [ND2] | 2.45 | A: NAI 601 [O2D] |

**Regulatory NADH interface: Hydrogen bonds**

| *No* | Structure1 | Dist (Å) | Structure2 |
|---|---|---|---|
| *1* | A: VAL 392[N] | 3.36 | C: NAI 603[O3D] |
| *2* | A: HIS 195[NE2] | 3.32 | C: NAI 603[O2D] |

**Supplementary table 3: Identifying hydrogen bond corresponding amino acids involved in NADH interfaces.** Figure A represent the closed GDH-GTP-NADH structure and figure B represent open GDH-GTP-NADH structure. The number of hydrogen bonds at NADH interfaces are same in both the condition however there are significantly changes in amino acids and corresponding hydrogen bond distances (Å). Moreover, in closed situation, catalytic NADH showing stronger interaction with NBD domain (all the hydrogen bond formation amino acid belongs to NBD domain).

A **GTP binding: Hydrogen bonds**

| *No* | Structure1 | Dist (Å) | Structure2 |
|---|---|---|---|
| *1* | A: ARG 265 [NH1] | 3.60 | A: GTP 602 [O1G] |
| *2* | A: ARG 265 [NH1] | 3.53 | A: GTP 602 [O2G] |
| *3* | A: TYR 262 [OH] | 2.35 | A: GTP 602 [O3G] |
| *4* | A: HIS 209 [NE2] | 2.36 | A: GTP 602 [O2B] |
| *5* | A: SER 213 [OG] | 2.75 | A: GTP 602 [O2] |
| *6* | A: GLU 292 [OE1] | 3.43 | A: GTP 602 [N1] |
| *7* | A: GLU 292 [OE2] | 2.72 | A: GTP 602 [N2] |

B **GTP binding: Hydrogen bonds**

| *No* | Structure1 | Dist (Å) | Structure2 |
|---|---|---|---|
| *1* | A: ARG 261 [NH2] | 2.78 | A: GTP 601 [03G] |
| *2* | A: TYR 262 [OH] | 2.84 | A: GTP 601 [O3G] |
| *3* | A: HIS 450 [NE2] | 3.23 | A: GTP 601 [O1B] |

C **GTP without NADH: Hydrogen bonds**

| *No* | Structure1 | Dist (Å) | Structure2 |
|---|---|---|---|
| *1* | A: SER 213 [OG] | 3.88 | A: GTP 601 [O1G] |
| *2* | A: ARG 261 [NE] | 2.74 | A: GTP 601 [O3G] |
| *3* | A: ARG 265 [NH2] | 3.20 | A: GTP 601 [O3G] |
| *4* | A: ARG 261 [NH2] | 2.71 | A: GTP 601 [O3G] |
| *5* | A: GLU 292 [OE2] | 3.13 | A: GTP 601 [N1] |

**Supplementary table 4: Identifying hydrogen bond corresponding amino acids involved in GTP interfaces.** Table A and B represent hydrogen bond with GTP in closed and open structure. Table C represent number of hydrogen bonds with GTP in GDH-GTP without NADH structure. So, in closed structure showing high volume of hydrogen bonds compare to open and GDH-GTP without NADH structure indicates stronger interaction of GTP with GDH in closed situation.

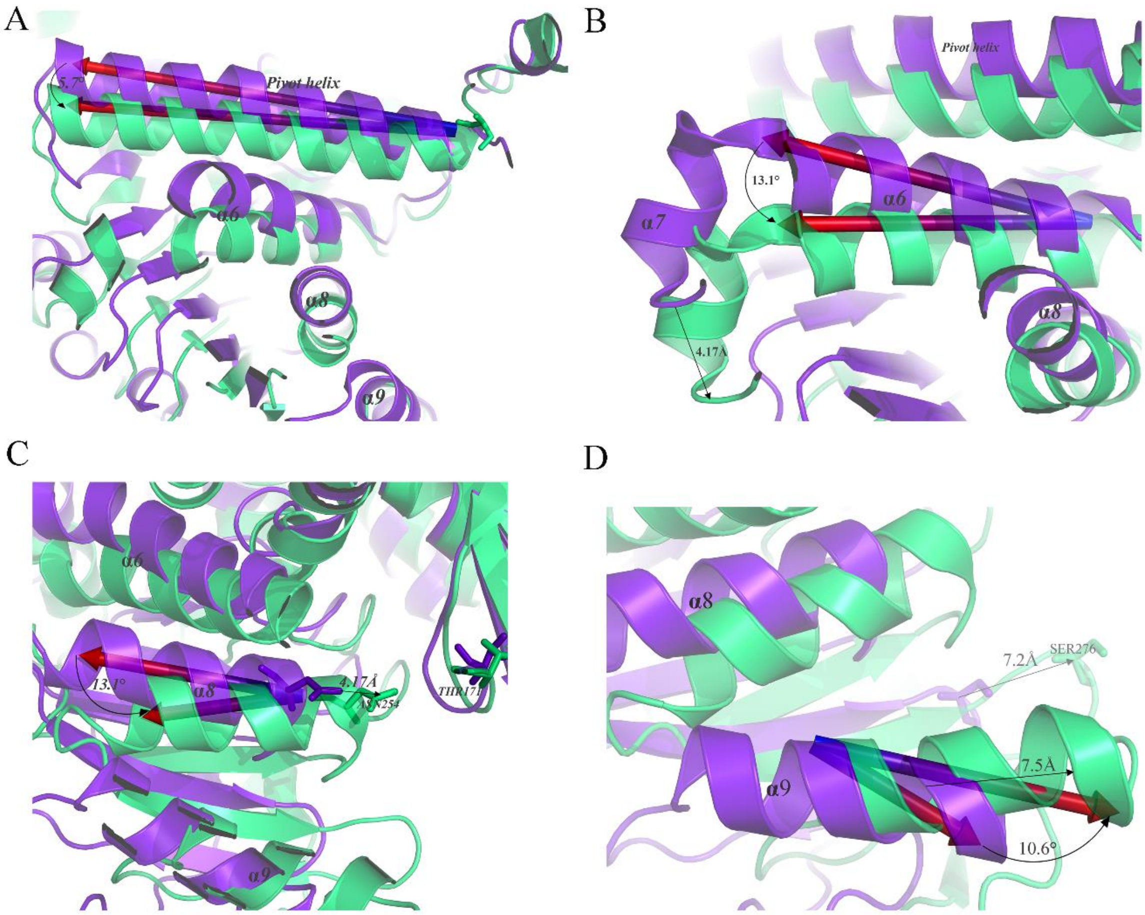


**Supplementary figure 4**: **Translation and rotation of superimposed helices** (open as purple and closed as lime green) those are situated at significantly high RMSD position (200 to 390) within protein. Pivotal and α6 helices is rotated (5.7$^{o}$ and 13.1$^{o}$) antilock wise however according to figure C and D, α8 and α9 helices is rotated (13.1$^{o}$ and 10.6$^{o}$) as well as translated (4.75Å and 7.5Å) towards catalytic site. This translation as well as rotation indicate the shifting (Ser276 is shifted approx. 7.2Å) of NBD domain towards catalytic site and closed the catalytic cleft. All the figure above is generated by UCSF Chimera.

| Number of mutations | Amino acid change in mature enzyme | "Sporadic"/ "Familial" | Exons | Positions in structure |
|---|---|---|---|---|
| 1 | SER ⟶ Cys at 213 | 0/1 | 6 and 7 | 4 Å of GTP |
| 12 | Arg ⟶ Cys at 217 | 3/9 | 6 and 7 | 4 Å of GTP |
| 2 | His ⟶ Thr at 258 | 2/0 | 6 and 7 | 4 Å of GTP |
| 2 | Arg ⟶ Thr at 261 | 2/0 | 6 and 7 | 4 Å of GTP |
| 1 | Arg ⟶ Ser at 261 | 0/1 | 6 and 7 | 4 Å of GTP |
| 1 | Tyr ⟶ Cys at 262 | 1/0 | 6 and 7 | 4 Å of GTP |
| 1 | Tyr ⟶ His at 262 | 1/0 | 6 and 7 | 4 Å of GTP |
| 13 | Arg ⟶ His at 265 | 2/11 | 6 and 7 | 4 Å of GTP |
| 1 | Arg ⟶ Cys at 265 | 1/0 | 6 and 7 | 4 Å of GTP |
| 1 | Leu ⟶ Val at 409 | 1/0 | 10,11 and 12 | Antenna |
| 2 | Asn ⟶ Tyr at 406 | 2/0 | 10,11 and 12 | Antenna |
| 1 | Phe ⟶ Leu at 436 | 1/0 | 10,11 and 12 | Antenna |
| 1 | Gln ⟶ Arg at 437 | 1/0 | 10,11 and 12 | Antenna |
| 4 | Gly ⟶ Val at 442 | 4/0 | 10,11 and 12 | Antenna |
| 3 | Gly ⟶ Asp at 442 | 3/0 | 10,11 and 12 | Antenna |
| 3 | Gly ⟶ Ser at 442 | 3/0 | 10,11 and 12 | Antenna |
| 1 | Gly ⟶ Cys at 442 | 1/0 | 10,11 and 12 | Antenna |
| 1 | Gly ⟶ Arg at 442 | 1/0 | 10,11 and 12 | Antenna |
| 25 | Ser ⟶ Leu at 441 | 24/1 | 10,11 and 12 | Antenna |
| 1 | Ala ⟶ Thr at 443 | 0/1 | 10,11 and 12 | Antenna |
| 3 | Ser ⟶ pro at 444 | 0/3 | 10,11 and 12 | Pivotal |
| 2 | Lys ⟶ Glu at 446 | 1/1 | 10,11 and 12 | Pivotal |
| 2 | His ⟶ Tyr at 450 | 1/1 | 10,11 and 12 | Pivotal |

**Supplementary table 5: Mutational hotspots region.** This table shows the location and frequency of hyperinsulinism–hyperammonemia syndrome (HI/HA) mutation in GDH. Out of total 84 cases, 66% is sporadic and 34% were Familial (Stanley et al., 2011). The table also indicate three mutational hotspot regions within protein, like GTP binding, Antenna and Pivotal helices region, however the highest number of mutations in HI/HA cashes has been observed at Leu441 whose position in the junction of pivotal and Antenna helices. Another position of mutation is His265 whose frequency also significantly higher in HI/HA cases.

| Structure ID | Hydrogen bond energy strength | Highest size rigid cluster | Participating helices (%, towards N or C terminal) within rigid cluster | Total No of Rigid Cluster | Independent DOF | No of hydrogen bond | Independent Hinge joint |
|---|---|---|---|---|---|---|---|
| 3jczA (Open) | 0.0 | 5274 | $\alpha1(91,N),\alpha2(93,C),\alpha3,\alpha4,\alpha5,\alpha6\&\alpha7(0.87,C)$ $\alpha8(67,C),\alpha9(92,N),\alpha10,\alpha11(93,N),\alpha15,\alpha16(90,N)$ | 1264 | 614 | 393 | 476 |
| | 0.5 | 4791 | $\alpha1(86,N),\alpha2(33,C),\alpha3,\alpha4,\alpha5,\alpha6\&\alpha7(87,C)$ $\alpha8(67,C),\alpha10,\alpha11(93,N),\alpha15,\alpha16(90,N)$ | 1448 | 663 | 349 | 506 |
| | 1 | 1867 | $\alpha4(20,N),\alpha5,\alpha6\&\alpha7(83,C),\alpha8(67,C),\alpha10,\alpha15,$ $\alpha16(67,N)$ | 2265 | 740 | 297 | 544 |
| | 1.5 | 1362 | $\alpha6\&\alpha7(45,N\&C),\alpha8(67,C),\alpha10,\alpha16(52,N)$ | 2721 | 836 | 251 | 567 |
| | 2 | 805 | $\alpha6\&\alpha7(45,N\&C),\alpha8(67,C)$ | 3009 | 909 | 220 | 585 |
| 3jd3A (Open) | 0.0 | 4809 | $\alpha1,\alpha2,\alpha3,\alpha4,\alpha6\&\alpha7(95,C),\alpha8,\alpha9(73,N),\alpha10(70,N)$ $\alpha11,\alpha15,\alpha16(90,N)$ | 1389 | 669 | 376 | 506 |
| | 0.5 | 2846 | $\alpha1(96,N),\alpha6\&\alpha7(95,C),\alpha8,\alpha9(73,N),\alpha10(70,N)$ $\alpha11(33,N),\alpha15,\alpha16(90,N)$ | 1935 | 725 | 324 | 536 |
| | 1 | 1471 | $\alpha6\&\alpha7(83,C),\alpha8,\alpha9(64,N),\alpha15(92,C)$ | 2479 | 809 | 280 | 570 |
| | 1.5 | 1025 | $\alpha6\&\alpha7(62,C),\alpha8,\alpha9(64,N)$ | 2690 | 895 | 236 | 588 |
| | 2.0 | 232 | $\alpha6\&\alpha7(38,C)$ | 3207 | 1017 | 190 | 618 |
| 3jd4A (Closed) | 0.0 | 5580 | $\alpha1(96,N),\alpha2,\alpha3(94,C),\alpha4,\alpha5,\alpha6\&\alpha7,\alpha8,\alpha9$ $\alpha10,\alpha11(95,N),\alpha15(67,C),\alpha16(92,N)$ | 1167 | 617 | 412 | 484 |
| | 0.5 | 5121 | $\alpha1(96,N),\alpha2,\alpha3(94,C),\alpha4,\alpha5,\alpha6\&\alpha7(96,C),\alpha8$ $\alpha9(75,N),\alpha10,\alpha11(84,N),\alpha15(66,C),\alpha16(92,N)$ | 1355 | 670 | 363 | 511 |
| | 1 | 4784 | $\alpha1(96,N),\alpha2,\alpha3(94,C),\alpha4,\alpha5,\alpha6\&\alpha7(96,C),\alpha8$ $\alpha9(75,N),\alpha10,\alpha11(84,N),\alpha15(66,C),\alpha16(92,N)$ | 1499 | 719 | 317 | 537 |
| | 1.5 | 1748 | $\alpha3(94,C),\alpha4,\alpha5(92,N),\alpha11(43,N)$ | 2478 | 803 | 270 | 551 |
| | 2 | 652 | $\alpha4(87,N)$ | 2908 | 903 | 226 | 580 |

**Supplementary table 6: Details table of rigidity based allosteric transmission using ProFlex/FIRST.** Hydrogen bond energy strength is considered from 0 to -2.0 as communication is started after hydrogen bond cut off 0.0 and has been stopped before cut-off -1.5 (supplementary figure). The table also indicated the position and percentage of helices those are involved with in rigid cluster at different hydrogen bond energy cut-off.

A

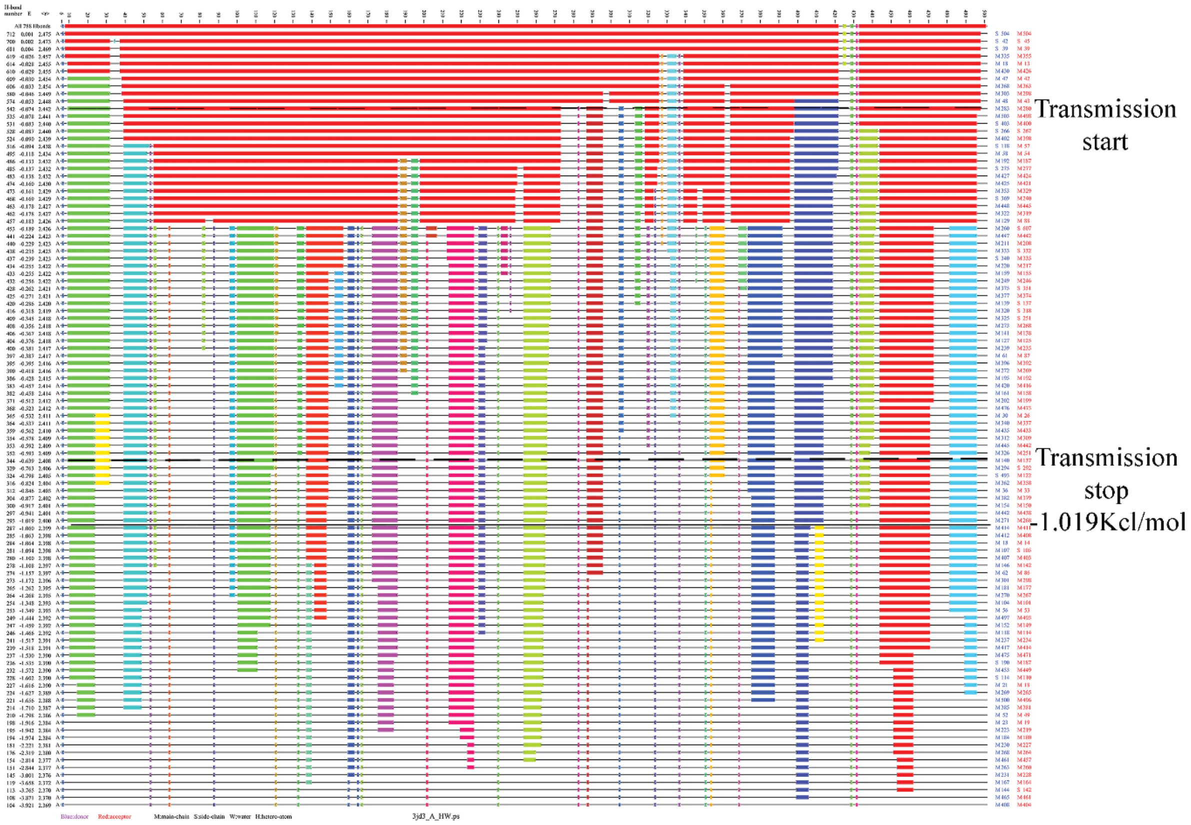

B

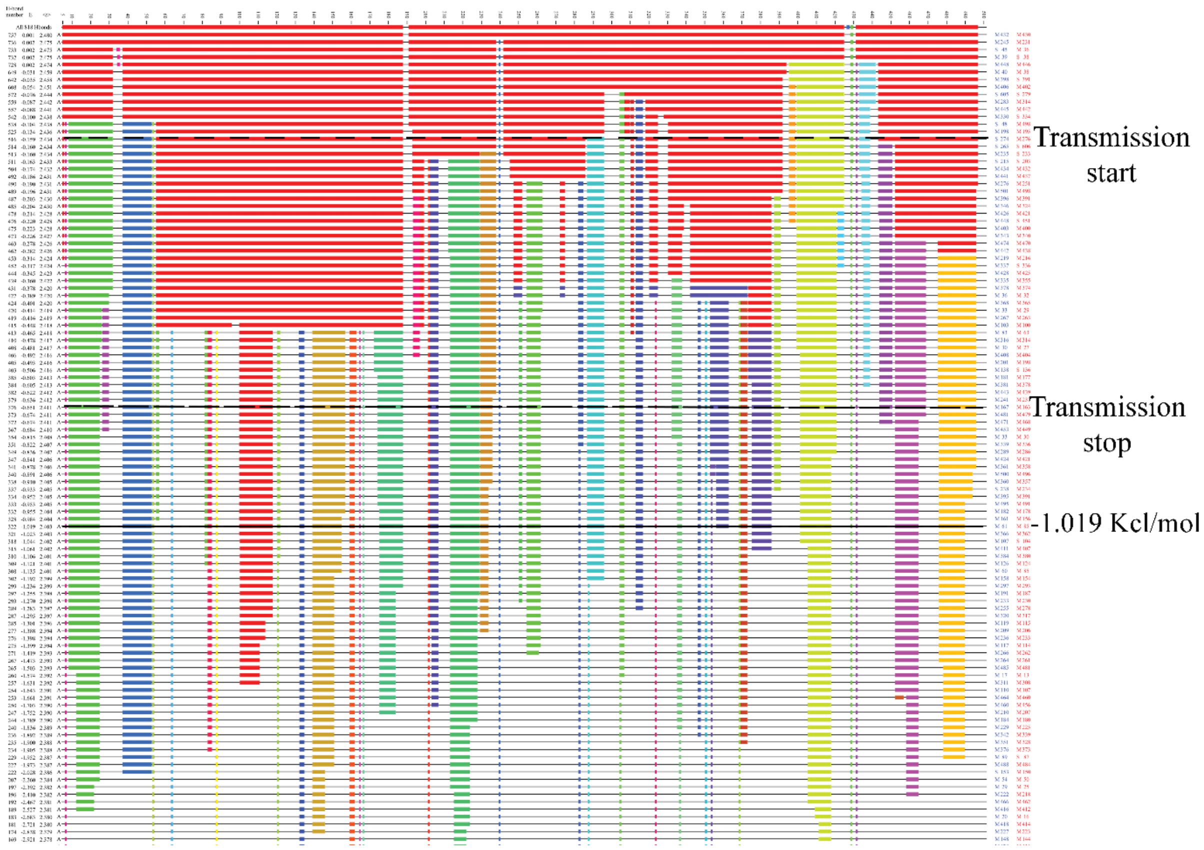


**Supplementary figure 5: Hydrogen bond dilution plot.** (A), (B) Dilution plot of GDH open (PDB ID 3jd3) and GDH closed (PDB ID 3jd4) using ProFlex / FIRST (Jacobs et al., 2001). Horizontal axis represents residue numbers and vertical axis represent hydrogen bond energy cut off (Kcl/mol). Protein flexibility shows by horizontal grey lines however solid colour lines indicate rigid cluster at different hydrogen bond energy cut off. Last two column (blue and red) represent hydrogen donor and acceptor respectively. Black dotted lines (upper and lower) indicates allosteric communication start and stop associated hydrogen bond energy cut off.

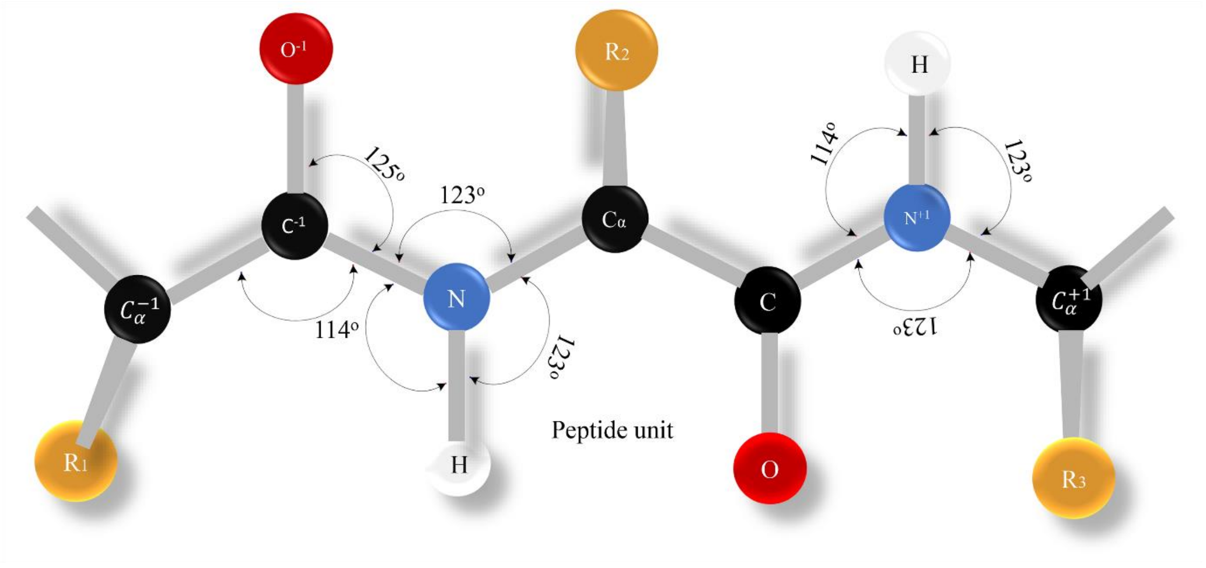

**Supplementary figure 6: Structural description of peptide chain geometry** i.e. amino acids in proteins (or polypeptides) are joined together by peptide bonds. $R_1$, $R_2$ and $R_3$ group represent the side chain of each amino acid.

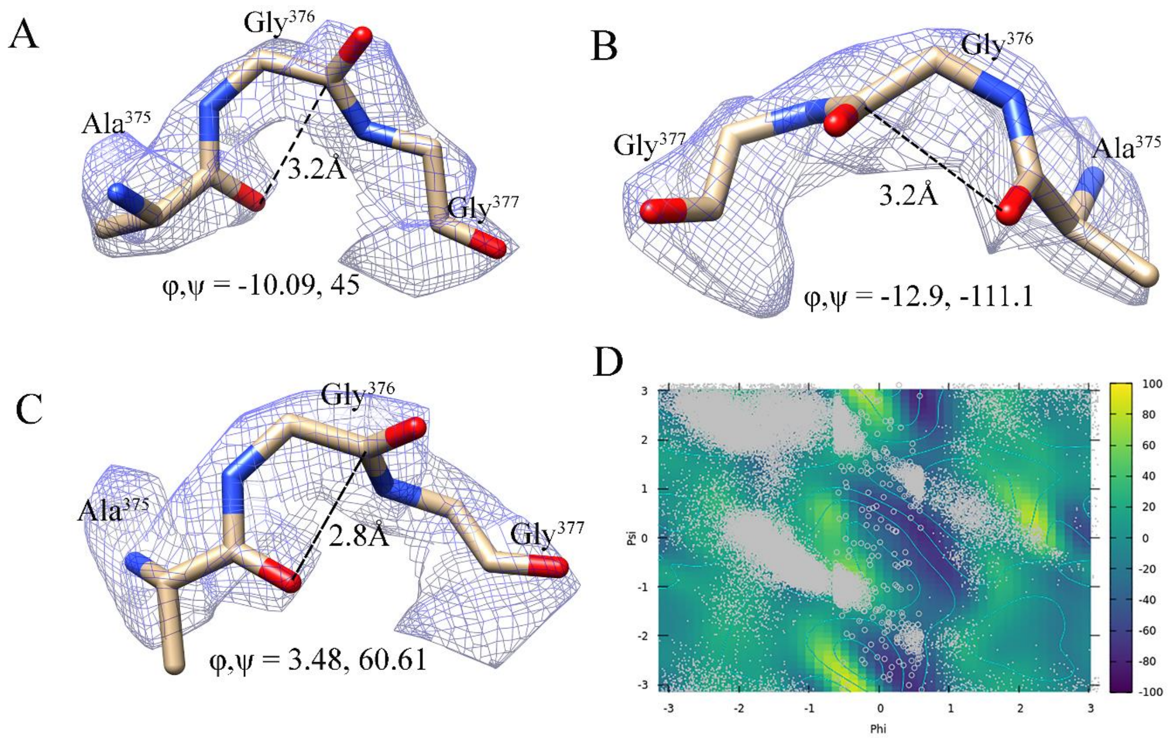


**Supplementary figure 7**: **Residue reliability and populated high-energy pass in transition upon mutation.** Figure (A), (B) and (C) showing the evidence of density map for a reliable residue at different time of trajectory (0.4ns, 9.3ns and 4ns) by adopting a confirmation at $-35 < \varphi < 35$. The residues Ala375, Gly376 and Gly377 from pdb snapshot at different time step (0.4ns, 9.3ns and 4ns) fitted with EM maps by chimera tool. Dotted black lines within figure is showing $O^{-1}$ …C distance corresponding to φ, ψ angles at different conformational states. (D) φ, ψ angles describing the local conformation changes of mountain pass residues at different time point trajectory after 10ns MD simulation upon mutant GDH structure (Gly376Asp). The zoomed circle indicates the position of transient residues at conformation $-35 < \varphi < 35$. Colour gradient also specifies the region of free energy landscape of Gly376 upon mutilation by Asp.

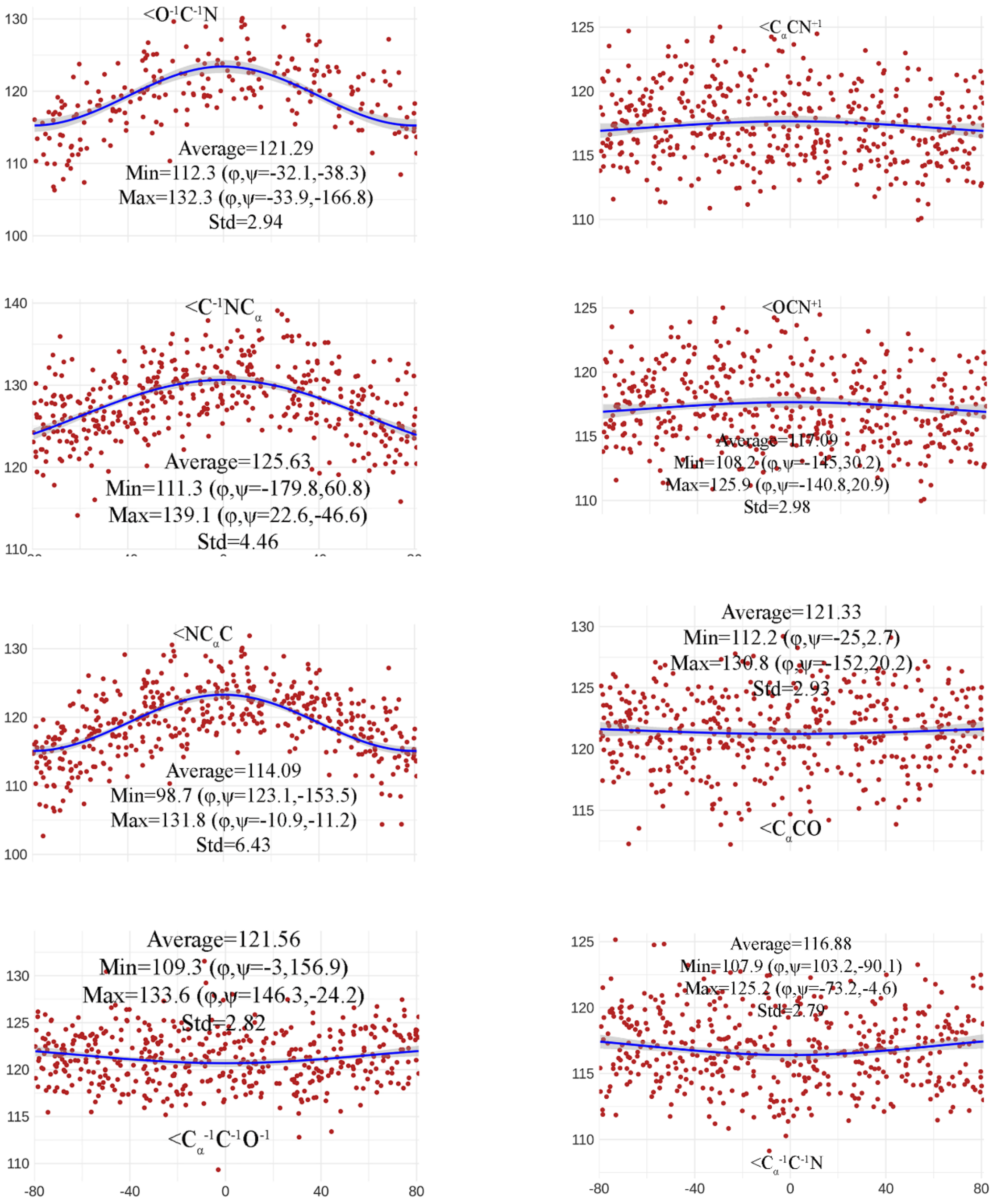


**Supplementary figure 8: Deformation of geometry with transition at high-energy pass.** Bond angle variations by obtaining the $\varphi \leq 0^{o}$ and $\varphi \geq 0^{o}$ transitions based on the MD trajectory data. Blue lines represent calculated fits of observed MD trajectory data using cosine equations to describe the systematic distortion of bond angles.

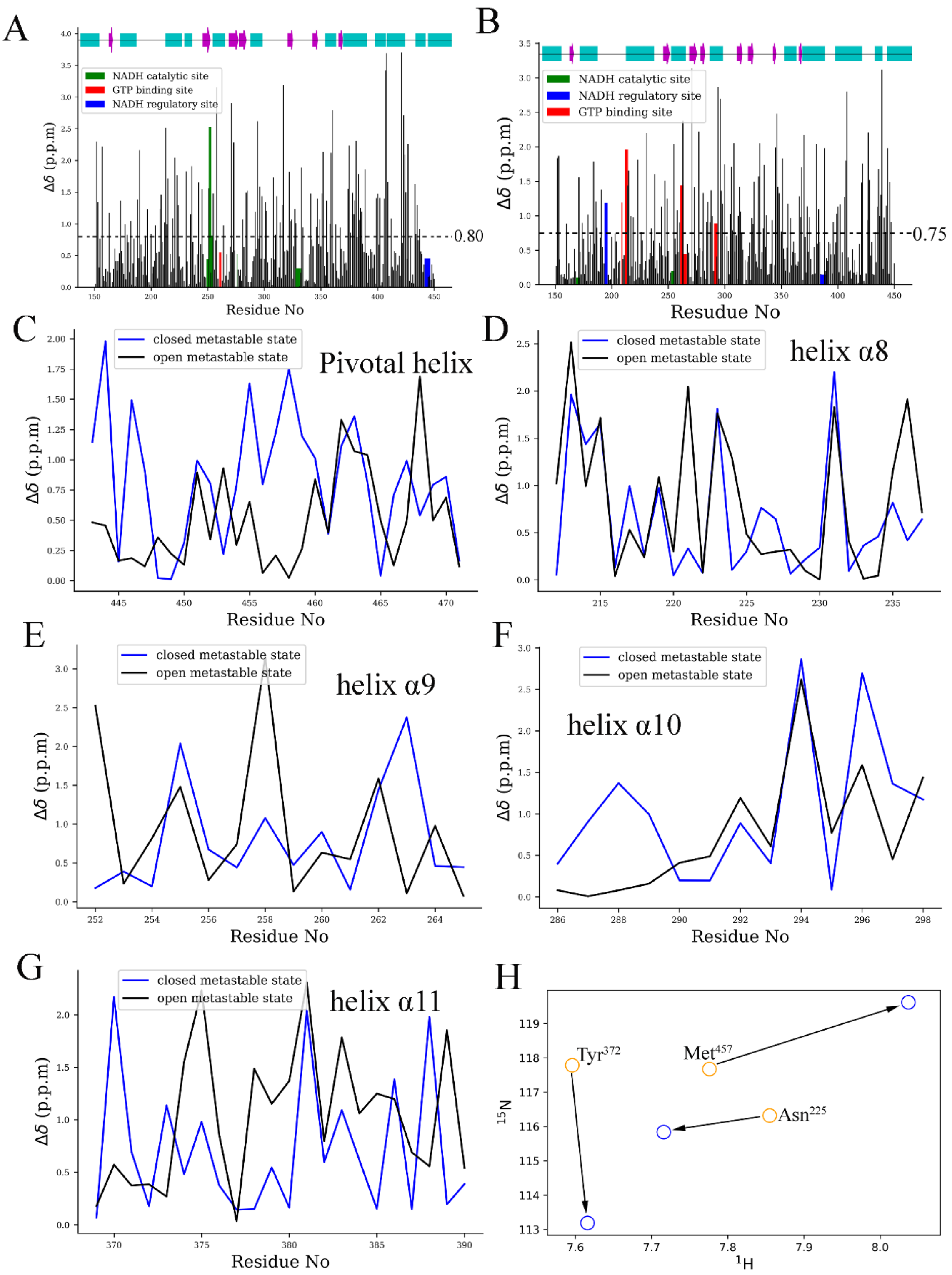


**Supplementary figure 9: Computation prediction of chemical shift.** Comparison of Chemical Shift between open and closed conformations was predicted by SHIFTX2 software (Han et al., 2011). (A) figure shows the difference of chemical shift in-between Apo (3jcz)

and NADH-GTP open (3jd3) structure. Colour area indicated the binding site of GTP and NADH. Top of the figure, rectangle and arrow with filled cyan and magenta colours mapped with alpha helices and beta sheets to indicate position of the residue. The dotted black line shows the average value of difference chemical shift (in ppm). (B) this figure also shows the difference chemical shift between Apo and NADH-GTP closed (3jd4) structures. Here the average chemical shift difference (in ppm) is slightly lower than former one. (C) figure shows 13CA chemical shift difference of pivotal helix where back and blue line represents NADH-GTP open and NADH-GTP closed structures. (D), (E), (F) and (G) also shows the 13CA chemical shift differences of α8, α9, α10 and α11 respectively. (H) 15N – 1H chemical shift differences of Tyr 372, Met 457 and Asn 225 indicates the loss of hydrogen bond in open structure (figure 2). All data were generated by SHIFTX2 (http://www.shiftx2.ca/) using experimental pH and temperature(K) available in PDB data (Han et al., 2011).

| Reside ID &NO | Frequency >=10 | Position |
|---|---|---|
| ALA-194 | 12 | Regulatory |
| ALA-341 | 31 | coenzyme binding domain |
| ALA-375 | 22 | Catalytic&376 |
| ALA-443 | 91 | Regulatory |
| ASN-254 | 20 | Catalytic |
| ASP-119 | 17 | -------- |
| GLU-25 | 42 | -------- |
| GLU-328 | 49 | Catalytic |
| GLU-36 | 19 | -------- |
| GLY-243 | 68 | coenzyme binding domain |
| GLY-350 | 53 | Catalytic |
| GLY-376 | 197 | Catalytic |
| GLY-377 | 19 | Catalytic |
| HIS-189 | 24 | Regulatory |
| HIS-209 | 31 | Regulatory |
| LEU-371 | 14 | coenzyme binding domain and alpha11 |
| LYS-329 | 28 | Catalytic |
| MET-169 | 63 | Catalytic |
| PHE-122 | 84 | --------- |
| PHE-304 | 10 | coenzyme binding domain |
| PRO-165 | 39 | Catalytic |
| PRO-167 | 40 | Catalytic |
| PRO-202 | 21 | Regulatory |
| PRO-240 | 32 | coenzyme binding domain |
| PRO-288 | 16 | coenzyme binding domain |
| PRO-354 | 19 | coenzyme binding domain |
| PRO-432 | 82 | Antenna |
| PRO-7 | 56 | N-terminal |
| PRO-88 | 15 | ------- |
| SER-327 | 33 | -------- |
| SER-393 | 41 | Regulatory |
| THR-34 | 18 | -------- |
| THR-37 | 36 | -------- |

| | | |
|---|---|---|
| THR-427 | 14 | Antenna |
| VAL-99 | 19 | Catalytic |

**Supplementary table 7: Frequency of amino acids in the high-energy transition region.** All the amino acid residues with frequency higher than 10 and fall into the region $-35 < \varphi < 35$ were taken after 10ns MD simulation upon GDH structure. Cyan colour indicates the frequency of Gly376 and its neighbourhood residues.

| **Reside ID &NO** | **Frequency >=10** | **Position** |
|---|---|---|
| ALA-336 | 10 | coenzyme binding domain |
| ALA-341 | 43 | coenzyme binding domain |
| ALA-434 | 25 | antenna |
| ARG-211 | 36 | GTP binding area |
| ARG-396 | 15 | regulatory |
| ASP-119 | 47 | ------ |
| ASP-138 | 12 | ------ |
| ASP-370 | 18 | coenzyme binding domain and alpha11 |
| ASP-376 | 168 | catalytic |
| GLU-173 | 25 | catalytic |
| GLU-25 | 60 | ------ |
| GLU-36 | 33 | ------ |
| GLU-38 | 13 | ------ |
| GLY-243 | 47 | coenzyme binding domain |
| GLY-350 | 147 | catalytic |
| GLY-377 | 16 | catalytic |
| GLY-422 | 34 | antenna |
| GLY-442 | 20 | regulatory |
| ILE-203 | 11 | regulatory |
| LEU-371 | 251 | coenzyme binding domain and alpha11 |
| LYS-329 | 26 | ------ |
| LYS-53 | 20 | ------ |
| MET-169 | 162 | catalytic |
| PHE-304 | 20 | coenzyme binding domain |
| PHE-9 | 18 | N-terminal |
| PRO-165 | 116 | catalytic |
| PRO-202 | 56 | regulatory |
| PRO-240 | 30 | coenzyme binding domain |
| PRO-354 | 17 | catalytic |
| PRO-429 | 43 | antenna |
| PRO-432 | 45 | antenna |
| PRO-7 | 40 | N-terminal |
| SER-276 | 58 | coenzyme binding domain |
| SER-279 | 20 | coenzyme binding domain |

| | | |
|---|---|---|
| SER-393 | 41 | regulatory |
| SER-83 | 19 | None |
| VAL-378 | 24 | catalytic |

**Supplementary table 8: Frequency of amino acids in the high-energy transition region upon mutation.** Amino acid residues with frequency more than 10 and falling in the high energy region -35 < φ < 35 were selected after 10ns MD simulation upon mutated GDH structure Gly376Asp. Gly376 and its neighbourhood residues were highlighted in cyan colour showing less frequency compared to previous table.
